# Septin-7 is indispensable for proper skeletal muscle architecture and function

**DOI:** 10.1101/2021.11.24.469846

**Authors:** Mónika Gönczi, Zsolt Ráduly, László Szabó, János Fodor, Andrea Telek, Nóra Dobrosi, Norbert Balogh, Péter Szentesi, Gréta Kis, Miklós Antal, György Trencsényi, Beatrix Dienes, László Csernoch

## Abstract

Today septins are considered as the fourth component of the cytoskeleton with the Septin-7 isoform playing a critical role in the formation of higher order structures. While its importance has already been confirmed in several intracellular processes of different organs, very little is known about its role in skeletal muscle. Here, using Septin-7 conditional knock-down mouse model, the C2C12 cell line, and enzymatically isolated adult muscle fibers the organization and localization of septin filaments is revealed, and an ontogenesis-dependent expression of Septin-7 is demonstrated. KD mice displayed a characteristic hunchback phenotype with skeletal deformities, reduction *in vivo* and *in vitro* force generation, and disorganized mitochondrial networks. Furthermore, knock-out of Septin-7 in C2C12 cells resulted in complete loss of cell division while KD cells provided evidence that Septin-7 is essential in proper myotube differentiation. These and the transient increase in Septin-7 expression following muscle injury demonstrate its vital contribution to muscle regeneration and development.

## Introduction

Proper contractile activation of the striated muscle requires precisely orchestrated machinery consisting of the T-tubule with the embedded dihydropyridine receptor (DHPR), the terminal cisternae of the sarcoplasmic reticulum (SR) with the ryanodine receptor (RyR) in its membrane, and mitochondria. Furthermore, appropriate localization of ion channels and proteins contributing to calcium homeostasis and downstream signaling pathways require a precisely organized structure and coordinated function of the cytoskeletal networks providing the connection between intracellular organelles (e.g. mitochondria) within the sarcomere, and the linkage between sarcomeres and surface membrane. Recent studies have suggested a role of septins [1] in these key cellular processes in cardiac and skeletal muscle function as well. However, muscle specific function of the different septin isoforms has not yet been completely explored.

Septins have been first described as regulators of cytokinesis and cell polarity in yeast, and since their discovery [2] their expression has been demonstrated in many other organisms, including mice and humans. Numerous studies imply the important role of septins in several intracellular processes as molecular scaffolds and diffusion barriers that control localization of membrane proteins. They are also involved in host–pathogen interactions during infections, in cell mobility [3], in apoptosis [4, 5] in endocytosis [6, 7], in determining cell shape [8], and even in mechano-transductional pathways [9–12]. Since septins interact with actin, microtubules, and membrane structures of cells and assemble into filaments, they became generally accepted as the fourth cytoskeletal component [12]. However, the molecular mechanism of these interactions is still the focus of intense investigation [13–15].

It is already known that these 30–65 kDa highly conserved GTP-binding proteins form hetero-oligomeric complexes and besides filaments, they polymerize into higher-order structures, sheets, rings, and cage-like formations contributing to biological processes [16, 17]. The number of septin isoforms encoded is extremely variable among different organisms. In humans 13 different isoforms were identified, which were classified into 4 groups (SEPTIN 2, 3, 6 and 7) on the basis of sequence homology [18], with a lone presence of Septin-7 in its group. Their ubiquitous [18–20] and/or tissue-specific expression influence the localization and function of cell surface proteins [9, 21]. The altered expression of septins has been linked to neurodegenerative or excretory system diseases, cardiovascular failures, immunological problems, or cancer [22].

Septin filaments are predominantly made up of hexamers and octamers of septins belonging to three or all four of homology groups, according to the cell type. All of these filaments include the Septin-7, which indicates the pivotal role of this ubiquitous protein in the formation of hetero-oligomeric complexes described to date [23–27]. The knockout of Septin-7 is embryonic lethal. Its absence resulted in the loss of other septin proteins in the oligomers [28]. Recently, the role of Septin-7 has been demonstrated in nervous and reproductive systems and its diverse functions in various neurological diseases (Alzheimer’s disease, schizophrenia, Neuropsychiatric systemic lupus erythematosus), in the development of cancer (glioma, papillary thyroid carcinoma and hepatocellular carcinoma). For a comprehensive review see Wang et al [29]. Furthermore, Septin-7 has been proposed as a novel regulator of neuronal Ca^2+^ homoeostasis [30, 31]. It has been revealed to downregulate the expression of the Orai and inositol-trisphosphate-receptor 3 (IP3R), subsequently affecting the cytosolic Ca^2+^ by which it can cause deficient flight ability in *Drosophila melanogaster* [31, 32]. Knockdown of Septin-7 resulted in myofibrillar disorganization and functionally reduced ventricular contractility and cardiac output in *Zebrafish* [33], caused an alteration of myosin heavy chain localization and disorganization of muscle fibers in somatic muscle [33], and significantly affected cytoskeleton dynamics in the ophthalmic artery [34]. To our knowledge, apart from these observations, no data is available on the function of Septin-7 in striated skeletal muscle.

Here, we give evidence on the crucial role of Septin-7 in skeletal muscle physiology, using the C2C12 cell line and a Septin-7 conditional knock-down mouse model established in our laboratory. We demonstrate an ontogenesis-dependent expression of Septin-7, its effect on the phenotype and the *in vivo* and *in vitro* force generation. Furthermore, we provide evidence that Septin-7 is essential in proper muscle development, in myotube differentiation and demonstrate its contribution to muscle regeneration.

## Results

### Skeletal muscle fibers express different septin isoforms

mRNA expression of representative members from all homology groups of septins has been detected in total lysates of human skeletal muscle by RT-PCR reactions on *m. quadriceps femoris* from amputated limbs (Figure 1A and Figure S1A). Most of the septin isoforms have been confirmed at the mRNA level in skeletal muscles originating from neonatal (5-day-old) C57BL/6J mice, as well. The expression pattern was similar in muscles of adult mice (4-month-old), although some of the isoforms (SEPTIN5-SEPTIN10) showed decreased intensity (Figure 1B). In parallel to the age-related alteration in mRNA expression of certain septin isoforms a differentiation dependent expression pattern of septins was revealed in C2C12 cells (Figure S1B). SEPTIN1 and SEPTIN3 were barely expressed, SEPTIN4 and SEPTIN7 expression showed gradual increase, while signals for other isoforms were rather uniform at different time points. SEPTIN12 and SEPTIN14-similarly to human and mouse muscle-were not detectable.

**Figure 1.**
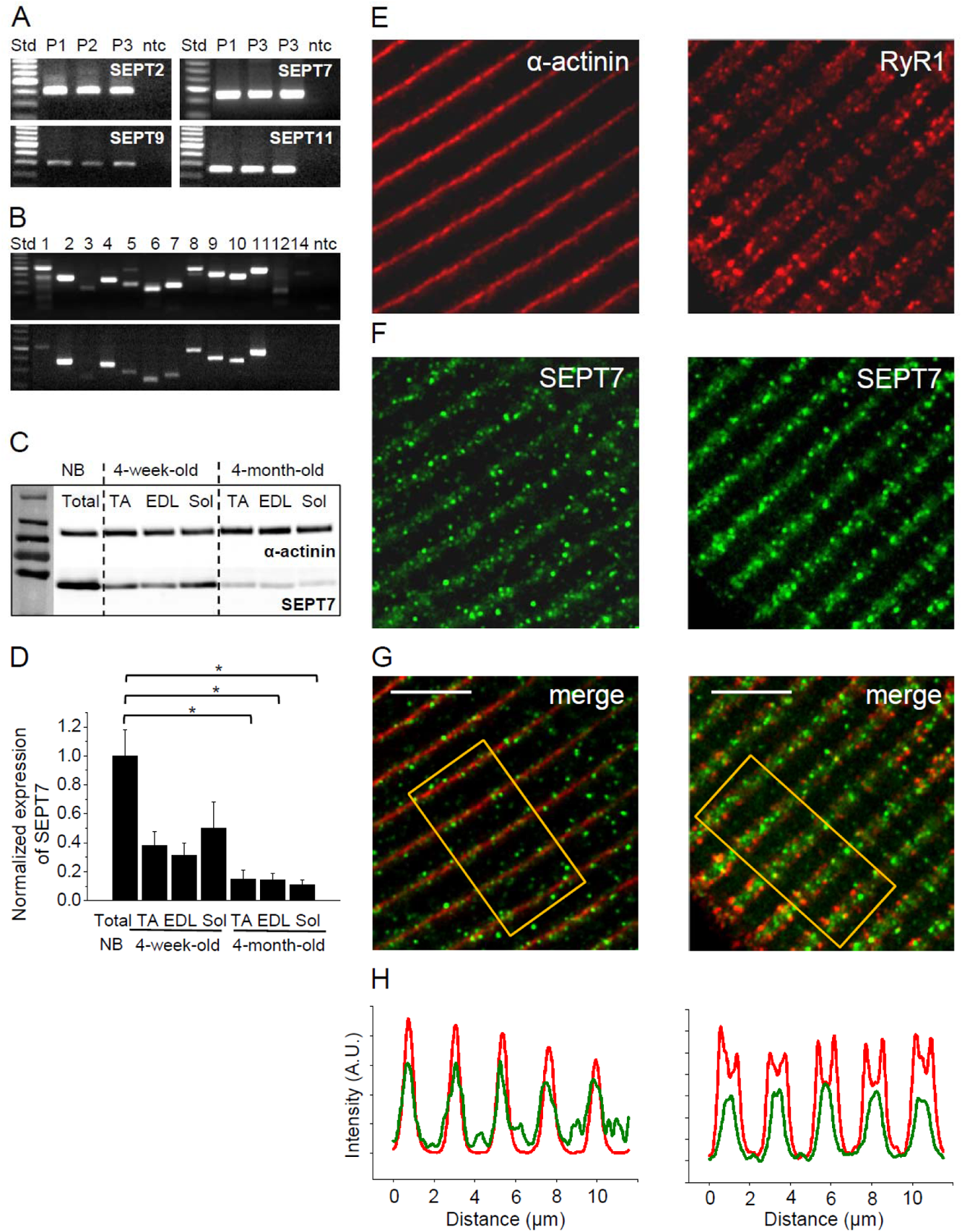
Septins are an integral part of the skeletal muscle cytoskeleton. **(A)** Agarose gel images showing the expression of septin isoforms at mRNA level in human skeletal muscle (m. *quadriceps femoris* from amputated limbs). Representative members from each homology group are shown. Independent samples from 3 patients were examined. **(B)** Expression of all septin isoforms at mRNA level in neonatal (upper panel) and adult (lower panel) mouse skeletal muscle. Non Template Control (ntc) samples contain nuclease free water instead of cDNA. **(C)** Differential expression of Septin-7 (SEPT7, 50 kDa) at protein level during development in different types of skeletal muscle in mice, α-actinin (110 kDa) was used as a normalizing control. Total skeletal muscle lysate of newborns (NB), TA: *m. tibialis anterior*, EDL: *m. extensor digitorum longus, and* Sol: *m. soleus*. **(D)** Pooled data of normalized Septin-7 expression during development. Representative data of 3 mice/age group. Data represent mean ± standard error of the mean, *p<0.05. Intracellular localization of Septin-7 (**F**) in adult skeletal muscle relative to α-actinin, and RyR1 (**E**) using immunofluorescence staining, and the merged images for the aforementioned proteins (**G**). Scale bar is 5 µm (**H**) Fluorescence intensity changes of Septin-7 (green) and α- actinin or RyR1 (red) along the fiber calculated from the rectangular area in panel G. See also Figure S1.

### Septin-7 shows ontogenesis dependent expression in skeletal muscle

Since the aforementioned alteration in its mRNA expression level during proliferation indicates a potential role in skeletal muscle development, Septin-7 expression was investigated in detail. Varying expression of Septin-7 has also been observed at the protein level during muscle development. Septin-7 protein expression declined gradually with age, as evidenced by comparing muscle samples from 4-week-old and 4-month-old mice to newborns. On the other hand, Septin-7 expression seems to be independent of muscle type, since signals were similar in different types of muscles (TA, EDL and Sol) in the selected time points (Figure 1C-1D). This ontogenesis dependent decline in the Septin-7 protein expression is consistent with our results at the mRNA level (Figure S1C). Septin-7 expression is remarkably elevated in proliferating cell lines and NB as compared to differentiating cultures and adult muscle samples, respectively (Figure S1D).

### Localization of Septin-7 on isolated skeletal muscle fibers

To assess the exact localization of Septin-7 within the skeletal muscle, enzymatically isolated FDB fibers from BL6 mice were subjected to specific immunolabeling. As the position of α-actinin and RyR1 is well defined in muscle fibers, they served as points of reference in these experiments (Figure 1E). Septin-7 was also visualized by immunocytochemistry (Figure 1F) and merged images were generated with α to determine the relative localization of Septin-7 (Figure 1G). Data acquired from confocal images showed that Septin-7 is located next to the Z-line similarly to α-actinin and Septin-7 signals were observed between the terminal cisternae characterized by RyR1 (Figure 1H).

### Skeletal muscle specific downregulation of Septin-7 results in an altered phenotype

To assess the function of Septin-7 in skeletal muscle, a mouse model with skeletal muscle specific knockdown of SEPTIN7 gene using the CRE/Lox system was generated (Figure 2A). Cre+ hemizygous mice following 3 months of tamoxifen feeding have been selected for the *in vivo* and *in vitro* experiments (Figure S1E). To estimate muscle-specific downregulation of Septin-7 protein expression, total lysates of *m. quadriceps femoris* and *m. pectoralis* were examined. Partial deletion resulted in a significant, and similar decrease of Septin-7 protein expression in both muscle types studied (to 59±8 and 63±8% as compared to Cre− mice, in *m. quadriceps* and *m. pectoralis*, respectively; Figure S1F-S1G) which correlated well with the approximately 50% deletion based on the three primer PCR method of gDNA that was achieved in the different skeletal muscle types (Figure 2B-2C). In addition, a pronounced spinal deformity was manifest in Cre+ mice in which the spine curved excessively outward, creating a hunchback (Figure 2A and Figure S2A-S2B). From CT images of three individual Cre+ and Cre− mice we determined the average angle of the 10^th^ thoracic vertebra (Figure S2C), and this parameter was significantly higher in Cre+ animals as compared to the Cre− littermates providing further evidence for the deformity of the vertebra (Figure S2D). As tamoxifen alone did not induce this deformity, it is likely to be the consequence of muscle atrophy and a weaker muscle tone. In line with the visible morphological changes, Cre+ mice had significantly smaller body weight starting from the age of the 7^th^ week (Figure 2D). The effect of tamoxifen treatment can be excluded since no significant difference was found between control BL6 and Cre− mice in terms of the body weight gain (Figure S3B).

**Figure 2.**
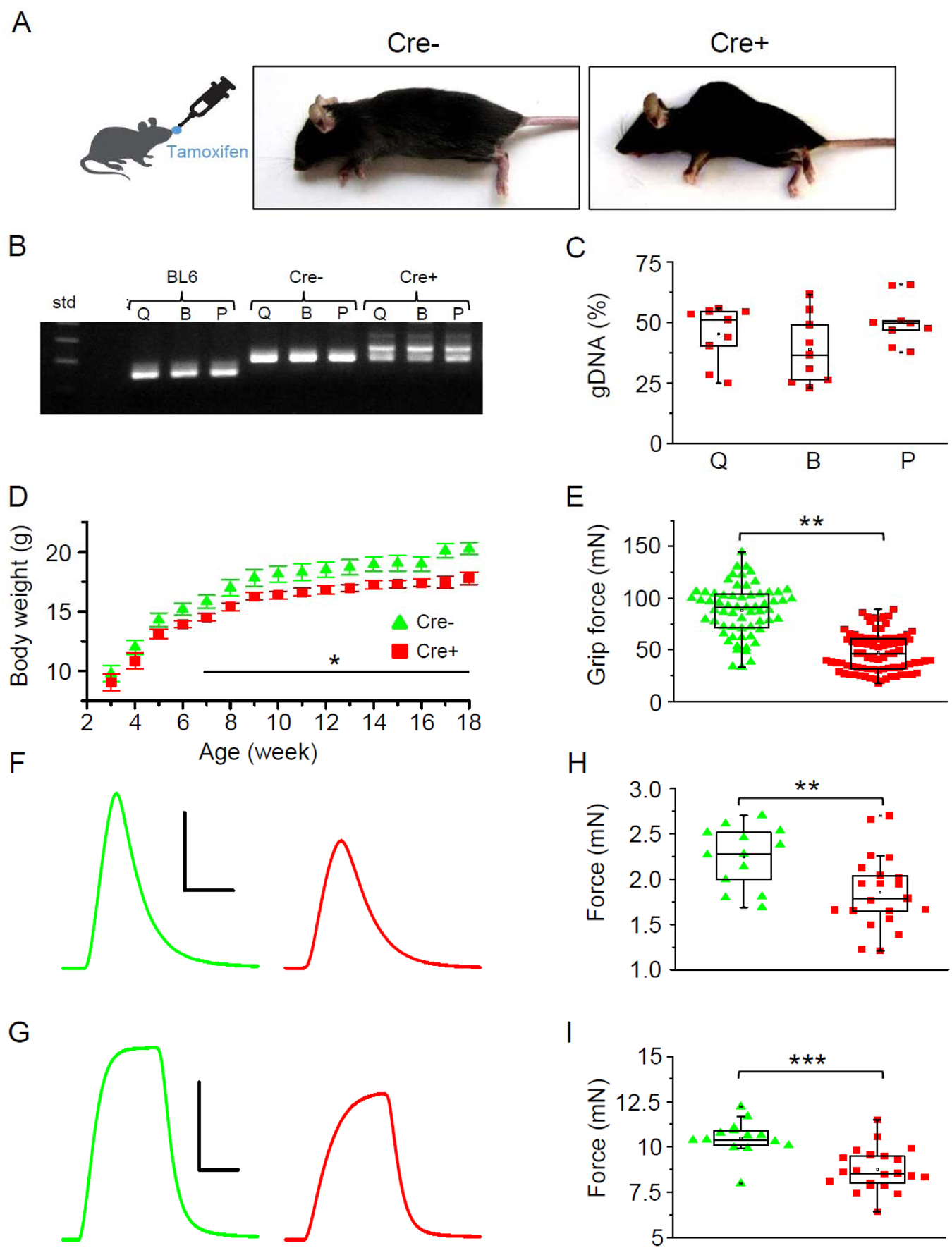
Skeletal muscle specific knock-down of Septin-7 resulted in a severe phenotype. *See also Figure S2; and Figure S3*. (**A**) Images of tamoxifen fed Cre− and Cre+ mice (both Septin-7^flox/flox^) at the age of 4 months. (**B**) Three primer PCR for detecting the partial deletion of the SEPTIN7 gene in different skeletal muscle types of Cre+ mice (Q: *m. quadriceps femoris*; B: *m. biceps femoris*; P: *m. pectoralis*) and the lack of deletion in samples prepared from BL6 control and Cre− mice. In samples originated from Cre− animals the floxed exon4, while in wild type tamoxifen fed mice the unmodified exon4 is demonstrated. (**C**) Pooled data of the percentage of exon4 deletion in different muscle types of Cre+ mice. 14 littermates (9 Cre+ and 5 Cre− were examined from 3 litters. Here and in all subsequent figures the rectangles in the box plots present the median and the 25 and 75 percentile values, while the error bars point to 1 and 99%. (**D**) Changes of body weight in Cre− (green triangle, n=11) and floxed Cre+ (red square, n=14) mice. Black solid line shows where the difference is statistically different (p<0.05). (**E**) Grip force in Cre− (n=4) and Cre+ mice (n=7). Representative twitch **(F)** and tetanic force **(G)** transients in EDL. Peak twitch **(H)** and tetanic force **(I)** in EDL from Cre− (n=7) and Cre+ (n=11) mice. Calibration in panel **F**: 1 mN and 50 ms; **G**: 5 mN and 100 ms. **p<0.01, ***p<0.001. See also Figure S2; and Figure S3.

### *In vivo* physical performance is impaired in Septin-7 knock-down mice

*In vivo* experiments have led to concordant observations, smaller body weight was accompanied by an impaired muscle performance. The average grip force was similar in control BL6 and Cre− mice (Figure S3C), while significantly smaller values were measured in Cre+ animals (Figure 2E). Voluntary running tests provided identical results. All parameters of running (distance, duration, average speed, maximal speed) were significantly smaller in Cre+ animals as compared either to BL6 or Cre− mice, while there was no difference between the parameters of running in BL6 and Cre− mice (Table S2).

### *In vitro* force is reduced in Septin-7 knock-down animals

The significant influence of Septin-7 reduction on muscle force parameters measured *in vivo* were also reflected in *in vitro* experiments. Both twitch and tetanic force decreased significantly in EDL (Figure 2F-2I) and in Sol muscle (Figure S3H-S3K) of Cre+ mice alike, as compared to the same muscles of Cre− littermates. This reduction in maximal force was the result of reduced Septin-7 content, since Tamoxifen treatment alone had no effect on the contractile parameters (Figure S3D-S3K). Interestingly, the kinetics as well as the fatigability of twitches and tetani did not change in Cre+ mice (Table S2). *In vivo* and *in vitro* force measurements thus suggest that Septin-7 fundamentally contributes to the normal skeletal muscle performance.

### Septin-7 is critical for proper cellular development and myotube differentiation

In proliferating C2C12 myoblasts the cellular distribution of Septin-7 (green) was detected along with the actin-network (red) using immunocytochemistry (Figure 3A), which revealed filamentous structure of Septin-7 and its co-localization with actin.

**Figure 3.**
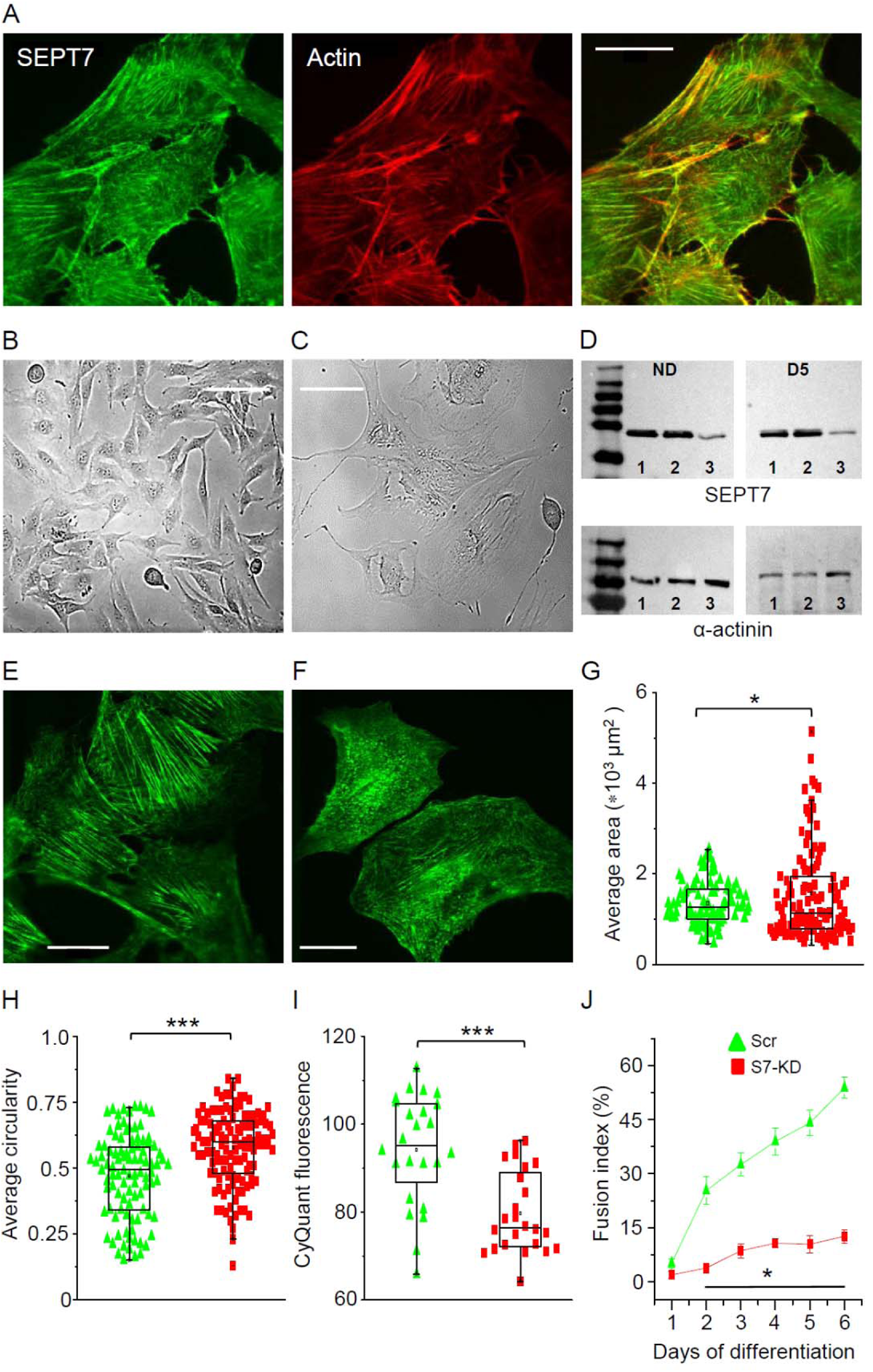
Septin-7 is critical for proper cellular development and myotube differentiation. (**A**) Confocal images of Septin-7 immunolabeling (green) and actin filaments (red) and their co-localization in control C2C12 cells. Calibration is 20 µm. Transmitted light images of control **(B)** and Septin-7 knockout (KO; **C**) cells demonstrating that complete KO of Septin-7 in C2C12 cells prevents appropriate cell proliferation. Scale bar for the control cells is 100 µm, while it is 50 µm for the KO cells. (**D**) Partial knock-down (KD) of Septin-7 expression at proliferation stage (non-differentiated, ND), and 5 days after differentiation induction (D5). 8µg of protein samples from absolute control (Ctrl), scrambled transfected (Scr), and S7-KD cells in each case were loaded to SDS-PAGE gel, and following electrophoresis and blot transfer into nitrocellulose membrane, α-actinin (110kDa) were probed with the appropriate primary antibodies. numbers 1, 2, and 3 at the bottom of the gels indicate Ctrl, Scr, and S7-KD samples, respectively. Immunolabeling of Septin-7 filaments in Scr **(E)**, and S7-KD cells **(F)** demonstrating altered filament structure, cell size, and shape. Scale bar is 20 µm. Quantification of changes in cell morphology, area **(G)** and circularity **(H)** in S7-KD cultures. Green triangles represent Scr, while red squares S7-KD cells. The number of cells investigated was 96 in Scr, and 121 in S7-KD cultures; *p<0.05, ***p<0.001; experiment was repeated twice (N=2). (**I**) Decreased proliferation 3 days after the plating assessed as CyQUANT fluorescence (n=24; N=3) and (**J**) suppressed differentiation of S7-KD cells determined by the calculation of fusion index during 6 days of investigation (n=20; N=2). Horizontal line in (**J**) shows where the difference between Scr and S7- KD cells was statistically significant (*p<0.05). See also Figure S4; and Figure S5.

The functional role of Septin-7 in C2C12 cells was evaluated using the CRISPR/Cas9 technique in control cells (Figure 3B). This approach led to abnormal cell size, in addition cell proliferation stopped (Figure 3C) providing an additional clue for the importance Septin-7 in cell division. Decreased expression of Septin-7 in C2C12 myoblasts was achieved using shRNA-mediated gene silencing. The suppressed protein level of knockdown (S7-KD) cells (Figure 3D) has been preserved along the differentiation process (Figure S4D) as compared to absolute control or scrambled shRNA-transfected cells. Having repeated the immunolabeling in scrambled (Figure 3E) and S7-KD cells (Figure 3F), strongly modified cell morphology and cytoplasmic distribution of Septin-7 was revealed. The characteristic filamentous structure of Septin-7 disappeared in most of the S7-KD cells, instead a fragmented, pointwise appearance of the protein was observed. In control and scrambled cultures Septin-7 and actin filaments were highly co-localized, while in S7-KD cells this spatial overlap was disrupted, although the actin structure within the cells seemed mostly unchanged (Figure S4A-S4C).

Changes in cell morphology were further analyzed, and significant differences in average cell area and circularity were found in S7-KD cells as compared to scrambled cultures (Figure 3G and 3H). These data express quantitatively the conspicuous visual observations, i.e. S7-KD cells became large in size and lost their processes, which resulted in a more rounded shape. At the same time, control and scrambled cells did not show any significant changes in respect to these parameters (Figure S4E, Figure S4F). Prominent changes in the intracellular architecture of Septin-7 in S7-KD cultured cells has been further proven by masking confocal images. Not only the significant changes in cell morphology are evident between the two cell types, but the remarkable alteration of filamentous structure is also well defined.

Myoblast proliferation capacity was also tested and significantly reduced proliferation was observed in Septin-7 modified cells (Figure 3I). Myotube formation was assessed by calculating the fusion index, the degree to which myogenic nuclei were found in multinucleated, terminally differentiated myotubes. From the second day of differentiation onwards increasing myotube formation was observed in control and scrambled cultures (Figure S4G), while only negligible cell fusion was detected in S7-KD cultures, even at 6^th^ day of the experiment (Figure 3J).

### *In vivo* knockdown of Septin-7 alters myofibrillar structure and mitochondrial parameters

Structural changes within the myofibrillar system triggered by the reduced Septin-7 expression was studied using electron microscopy (EM) on TA muscle. Representative EM images taken on transversal muscle sections of Cre− animals (Figure 4A) demonstrated the presence of myofibrils of a single fiber, which are all well demarcated with a visible sarcoplasmic reticulum. In samples of Cre+ mice (Figure 4B), however, separation of myofibrils was less obvious. After identifying all individual myofibrils within an average area of the actual visual field, average area, perimeter of the individual myofibrils, and the average number of myofibrils have been estimated within a given unit of area (1 µm^2^) of the actual visual field. As Figure 4C demonstrates, both the average area and the perimeter of myofibrils from Cre+ animals were significantly smaller than the corresponding parameters of myofibrils from Cre− mice. While the number of myofibrils within a given area of the visual field (Figure 4D) was increased significantly in samples originating from Cre+ mice as compared to sections of Cre− animals. In myofibrils from Cre+ mice all the aforementioned parameters were also significantly different from the control BL6 samples. On the other hand, there was no significant difference between the corresponding parameters of myofibrils originating from BL6 and Cre− animals (Figure S6C-S6D), suggesting that the markedly reduced size and increased number of the individual myofibrils per unit area in Cre+ mice are attributable to the modified Septin-7 expression in their skeletal muscle.

**Figure 4.**
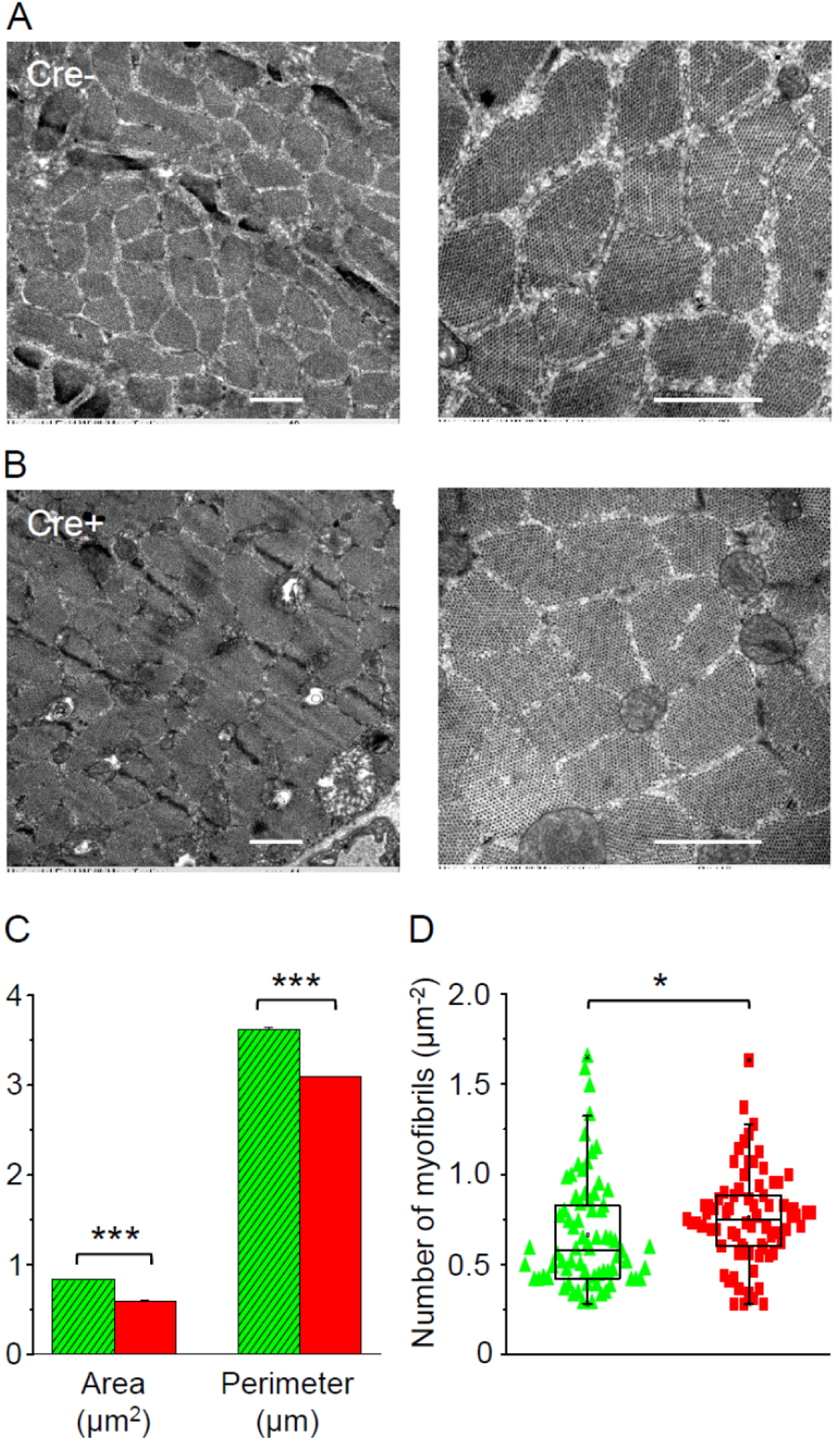
*In vivo* knockdown of Septin-7 alters myofibrillar structure. (**A-B**) Representative EM images of myofibrils from cross sectional samples of TA muscles from Cre− and Cre+ mice at smaller (left) and larger (right) magnification where scale bars represent 1 μm. (**C-D**) Area and perimeter of the individual myofibrils and the number of myofibrils within 1 µm^2^ of an appropriate visual field were determined from EM images. Here and in all subsequent figures green columns represent data from Cre−, while red columns from Cre+ animals (average±SE). The total number of myofibrils counted for area and perimeter were 3012 and 3174 in Cre− and Cre+ mice, respectively, while the number of visual fields examined was 72 and 74, for calculating the number of myofibrils (*p<0.05; ***p<0.001). See also Figure S6.

The aforementioned parameters have also been calculated for the mitochondria using similar transversal sections of TA muscles dissected from the different animal groups. In Figure 5A and 5B representative images on muscle sections from Cre− and Cre+ mice are presented, respectively. The calculated average area and perimeter of the individual mitochondria were found to be significantly reduced in Cre+ samples as compared to the mitochondrial data of either muscles from Cre− animals (Figure 5C) or to the samples of control BL6 mice. The number of mitochondria per unit area within the selected visual fields, however, was significantly increased in samples from Cre+ mice in comparison with the muscles of Cre− or BL6 mice, as shown in Figure 5D. Mitochondria-related parameters were not different between BL6 control and Cre− groups (Figure S7C-S7D).

**Figure 5.**
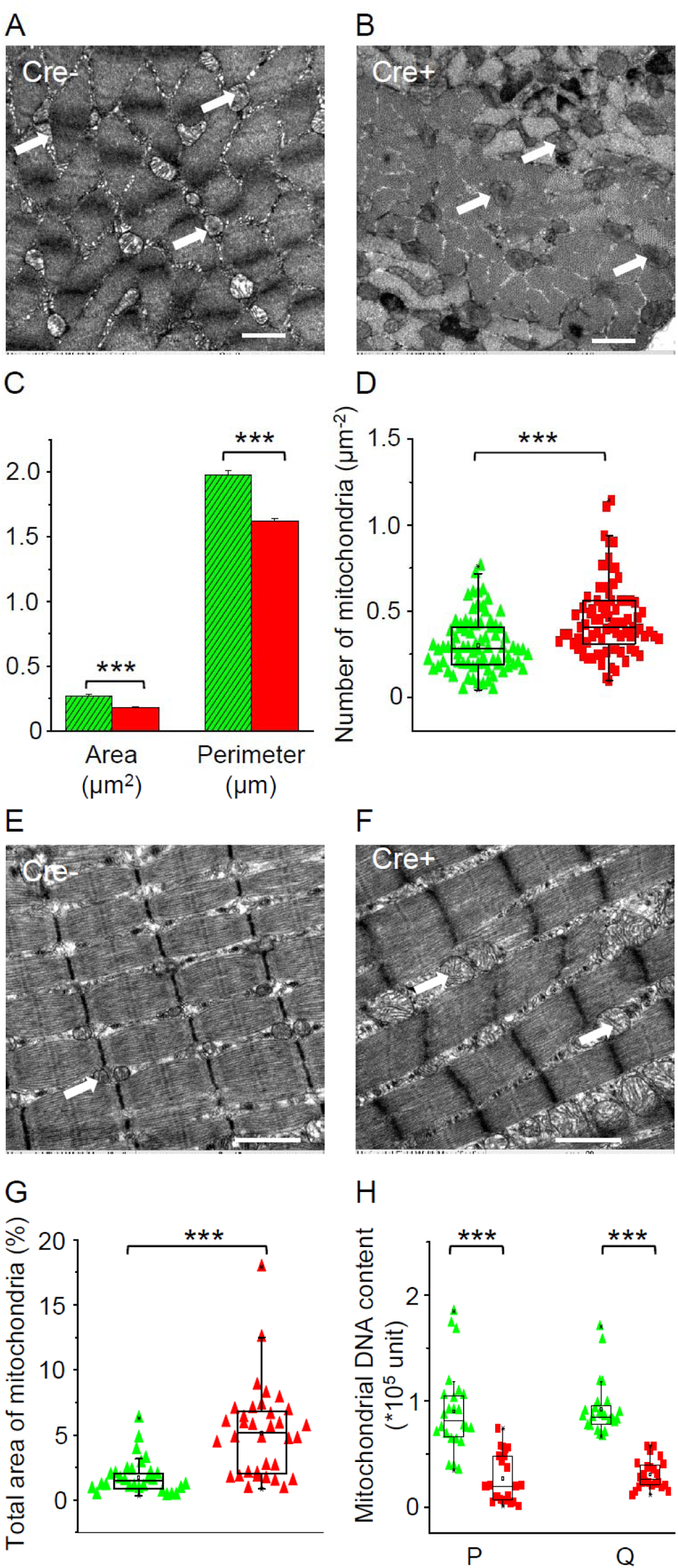
*In vivo* knock-down of Septin-7 alters mitochondrial parameters. (**A-B**) Representative EM images of myofibrils from cross sectional samples of TA muscles of Cre− and Cre+ mice. (**C-D**) Area and perimeter of the individual mitochondria and the number of mitochondria within 1 µm^2^ of an appropriate visual field were determined from EM images. The total number of myofibrils counted for area and perimeter was 1589 and 2105 in Cre− and Cre+, respectively, while the number of visual fields examined was 81 and 83 for calculating the number of mitochondria (***p<0.001). (**E-F**) Representative EM images of myofibrils and mitochondria from longitudinal sections of TA muscle samples originating from Cre− and Cre+ mice. (**G**) The total area of mitochondria within the actual images were determined and given as a percentage of the total area of the visual field. Number of visual fields investigated were 29 and 34, respectively, ***p<0.001. Scale bars are equal to 1 µm in all images. (**H**) Mitochondrial DNA content was determined from *m. pectoralis* (P) and *m. quadriceps femoris* (Q) of Cre− and Cre+ mice using specific qPCR primers. Number of samples were 8, while in each case experimental triplicate was performed, ***p<0.001. See also Figure S7.

Transmission electron microscopic analysis of longitudinal sections from the different animal groups revealed normal myofibrillar structure (sarcomere length, triad composition) in Cre− samples, while occurrence of large mitochondrial networks have been identified in most images taken from Cre+ muscles, as demonstrated by representative images in Figure 5E and 5F, respectively. The total area of mitochondria as a percentage of the total area of the corresponding visual field has also been determined, and this parameter in samples from Cre+ mice greatly exceeded the values from both the sections of Cre− and the BL6 mice), respectively (Figure 5G). There was no significant difference between these parameters of muscle sections originating from BL6 and Cre− animals (Figure S7E).

To illustrate better the lack of change in the area and perimeter of myofibrils/mitochondria in Cre− and BL6 control samples, statistical distribution of the datasets for all parameters were calculated and plotted as histograms. In Figure S6A-6B and Figure S7A-7B area and perimeter of myofibrils and mitochondria, respectively, show no or very slight shift between samples from control and Cre− mice.

Mitochondrial DNA content in different muscles, *m. pectoralis* and *m. quadriceps*, were also determined. Normalized mitochondrial (16S) RNA amount was found to be severely reduced in muscles from Cre+ mice as compared to data of Cre− mice in both examined muscles, respectively (Figure 5H).

### Septin-7 expression is modified during skeletal muscle regeneration

Previous data (shown in Figure 1C-1D and Figure S1C-S1D) suggested that Septin-7 has a prominent role in muscle development and structural assembly of newborn animals and in proliferating C2C12 cells, so its contribution to muscle regeneration was analyzed. Mild muscle injury in TA muscles was induced by *in vivo* BaCl_2_ injection in mice. At two different time points cryosections were prepared from the injected, and the contralateral, non-injected control muscles and were subjected to HE-staining. Figure S8 presents images from non-injected (A) and injected muscles (B) (as indicated) 14 days after the treatment. Satellite cell-coupled Pax7 expression was observed during regeneration. More intensive PAX7 positive signal and regeneration-induced, central nuclei in the myofibrils were detected in injected muscles (see magnified region of Figure S8A and S8B) in Figure 6B as compared to non-injected controls (Figure 6A) using DAB reaction. As demonstrated in Figure 6D and Figure 6F, the level of Pax7 expression was found to be significantly higher in the injected than in the non-injected muscles. Septin-7 protein expression was also monitored for two weeks during regeneration (Figure 6C) and western blot analysis revealed pronounced overexpression at each time point investigated (Figure 6E). Similar upregulation of Septin-7 and Pax7 detected upon BaCl_2_-induced muscle injury suggests that cytoplasmic septins have an essential role in regeneration via regulating proliferation and differentiation of newly formed muscle cells.

**Figure 6.**
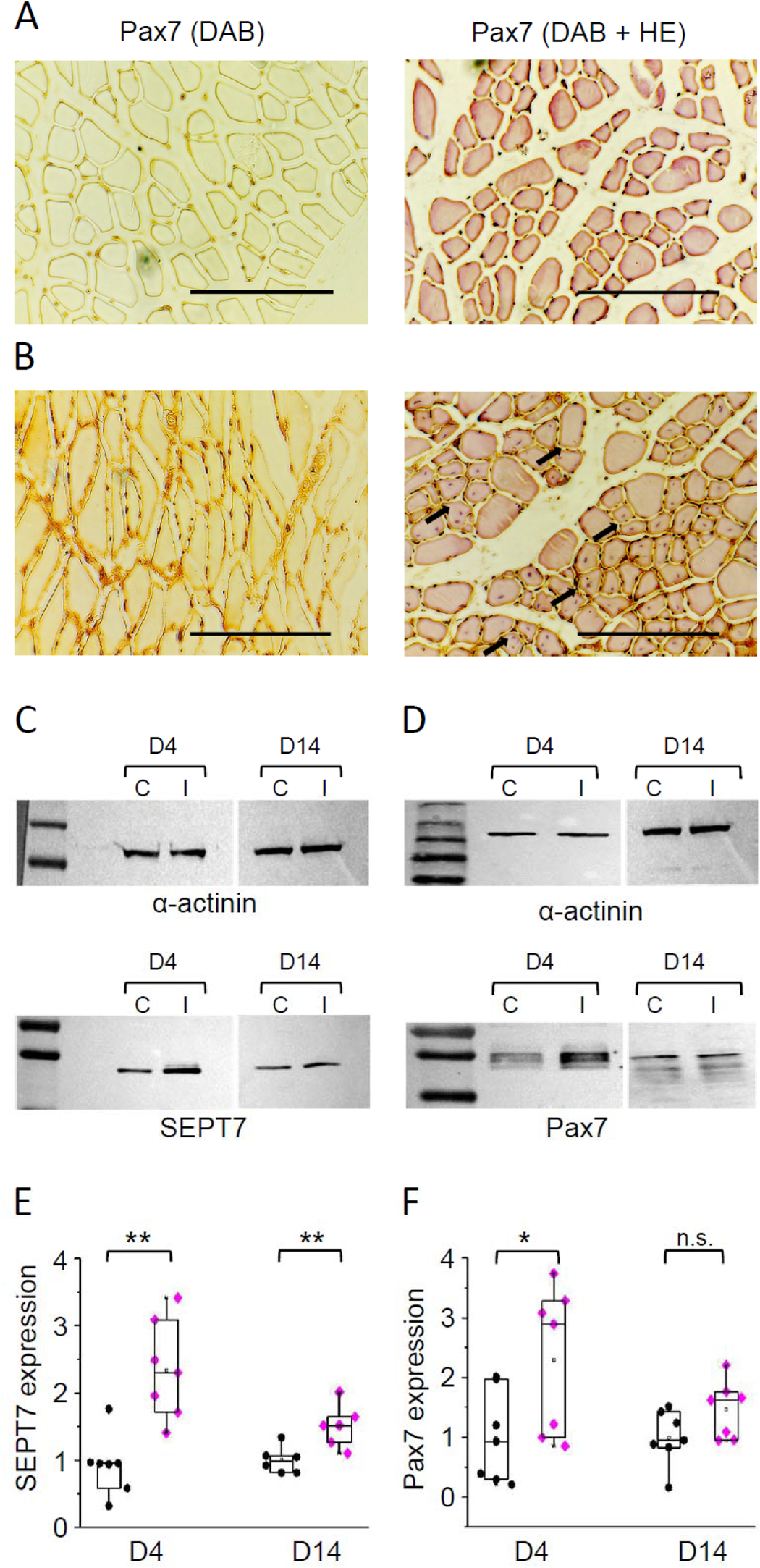
Septin-7 and muscle regeneration. Representative histological images from non-injected (A) and BaCl_2_-injected (B) TA muscles of BL6 mice 14 days following the muscle injury. The 6 µm cryosections were subjected to Pax7-specific immunostaining either alone (left panels) or together with HE-staining (right panels), latter represents regenerating myofibrils with central nuclei in BaCl_2_-injected samples (some indicated by arrows). Scale bars are equal to 100 µm. **(C-D)** Protein samples were prepared from control (C) and injected (I) muscles at appropriate time points (day 4-D4 and day 14-D14). Septin-7 and Pax7 protein expression was determined in each sample pair and α-actinin was used as normalizing control. Normalized Septin-7 **(E)** and Pax7 **(F)** expression level was determined in control (black circles) and injected muscles (magenta diamonds), and plotted individually at each time point of investigation. Each point represents individual data, *p<0.05; **p<0.01. See also Figure S8.

## Discussion

### Septin filaments are integral part of the skeletal muscle cytoskeleton

Specific members of different septin isoforms are present from yeasts to mammals during the phylogenesis (see review [1]). They have been reported to interact with other cytoskeletal elements, with the plasma membrane, and furthermore, to be involved in a number of signaling pathways [35–39]. In this study we demonstrate for the first time the presence of different septin isoforms and the contribution of Septin-7 to skeletal muscle physiology.

### Expression of septin isoforms in skeletal muscle

As septins are known to be the fourth component of the cytoskeleton, they could participate in the skeletal muscle architecture and function as well. Based on their mRNA expression profile in mouse and human skeletal muscle samples, the presence of at least one essential member of each homology group indicates the possibility of build-up of the previously described hetero-oligomeric structure [23–27]. These results predict the presence of a higher-order oligomeric structure of septins, though further structural investigations are needed.

Septin-7 showed a differential, declining expression with age both in fast and slow twitch skeletal muscles, which may emphasize its significance in muscle development. In line with the observation, that genetic deletion of SEPTIN7, SEPTIN9, or SEPTIN11 resulted in embryonic lethality, reflecting their essential role in early, intrauterine development [40–42]. Considering that SEPTIN7 is the only member of the SEPTIN7 homology group, its loss is not expected to be compensated for in the oligomers. Furthermore, the conditional SEPTIN7 KD (Cre+) mice showed altered phenotype and produced decreased muscle force. This raises two questions: (i) are septins essential in the early development of skeletal muscle, and (ii) do they lose their significance with aging?

Connection between septins and other cytoskeletal proteins has a prominent role in the conversion of mechanical inputs into biochemical signals. Septins have been shown to colocalize with actin filaments within stress fibers associated with focal adhesion complexes, as well as perinuclear actin. Septins participate in mechano-transduction by promoting the formation of contractile actomyosin networks, and the recruitment of myosin to actin in cancer-associated fibroblasts, in mammalian epithelial cells, and mouse cardiac epithelial cells (see the review [11]). In accordance with the above, we have demonstrated the co-localization of Septin-7 and actin filaments in control C2C12 cells, and furthermore, without Septin-7 these cells were not able to proliferate. Knocking down Septin-7 in C2C12 cells resulted in altered filament structure, cell size and shape as well. This is in line with the observation that during embryogenesis, interaction of Septin-2, 6, 7, and 9 with the cytosolic elements has an essential role in the regulation of cardiac functions [43].

### Suppression of septin expression is skeletal muscle results in severe skeletal deformities

Although septins were considered as key players of cell division based on their evolutionary conserved function in cytokinesis, they are now associated with a variety of other processes. Downregulation of the ubiquitous Septin-7 has been implicated in several pathological events including abnormal morphology and immature sperm [44]; altered glucose uptake in the insulin-sensitive podocytes [45]; inefficient microvascular angiogenesis, actomyosin organizations, and directional migration in primary mouse cardiac endothelial cells [46]. Septin-7 has been suggested as novel regulator of neuronal calcium homeostasis, since in resting neurons suppressed dSEPTIN7 (homologue of human SEPTIN 7 in *Drosophila*) resulted in Ca^2+^ store-independent opening of dOrai, a calcium release activated calcium channel [31]. Also, Septin-7 deficiency alters the morphology of mature neurons [32,47–49], leading to neurological disorders such as Alzheimer’s disease, schizophrenia, neuropsychiatric lupus erythematosus.

In Septin-7 modified Cre+ mice, generated in our laboratory, spinal structural deformity, similar to the human Scheuermann’s kyphosis [50] affecting the thoracic or thoracolumbar spine was observed. The muscle atrophy and decreased muscle tone observed in Cre+ animals are assumed to be responsible for this vertebral defect, since tamoxifen treatment alone did not cause similar malformation. In itself, tamoxifen had no effect on muscle structure or body weight. It should be noted, however, that such effects of tamoxifen were administered to mice with impaired muscle function, i.e. to mdx mice (mouse model of Duchenne muscular dystrophy) [51], to FKRPP448L mutant mice with dystroglycanopathy [52], and to Mtm1 knockout mice (murine model of myotubular myopathy) [53]. The latter is the only study in which wild type control mice have also been used.

### Role of septins in determining cell proliferation and cell morphology

Complete knockout of Septin-7 was not possible in cultured C2C12 cells, however, reduced Septin-7 expression significantly decreased proliferation rate and inhibited myotube formation. This is in line with previous observations that septins are essential regulators of cell division from yeast to *Drosophila* [54–59], and in CHO cells [60]. In epithelial cells septin filaments are involved in chromosome movement and spindle elongation during mitosis [61, 62], and even cytokinesis of T cells requires the presence of septins [12, 63].

Furthermore, reduced Septin-7 expression resulted in large and nearly circular C2C12 myoblast cells, with a deformed intracellular filamentous septin structure. This correlates with the observation that Septin-7 determines cell morphology in different cell types, like dorsal root ganglia neurons in the nervous system [64, 65], or in mouse primary fibroblasts [66]. Finally, involvement of Septin-7 as a target gene of miR-127-3p has been revealed in the regulation of C2C12 proliferation [67]. These observations together with our data suggest that cytoskeletal septin organization has a pivotal role in determining cell morphology, proliferation, and differentiation of skeletal muscle cells.

### Role of septins in mitochondrial dynamics

Our results suggest the involvement of septins in mitochondrial dynamics in skeletal muscles from mice, since mitochondrial area and number were altered following the conditional knockdown of Septin-7 expression. Furthermore, long mitochondrial networks were found within these fibers. However, mitochondrial DNA content was reduced, suggesting impaired mitochondrial function in line with Afshan and Czajka [68] in Cre+ mice.

The cytoskeleton has been shown to alter mitochondrial movement and distribution in highly polarized cells and to play a role in mitochondrial dynamics (see the review [69]). In particular, in septin depleted cells elongated mitochondria developed due to decreased fission rather than to a defective mitochondrial fusion [70]. Depletion of Septin-2 has been shown to influence mitochondrial morphology in HeLa cells [70], and in *Caenorhabditis elegans* [70], suggesting that the role of septins in mitochondrial dynamics is evolutionarily conserved. The contribution of mitochondria-associated septin cages is also described; e.g. Septin-7 co-localization with mitochondria was shown to affect the proliferation of *Shigella flexneri* [71], while disrupted mitochondrial morphology was detected in the ciliate *Tetrahymena thermophila* following modification of septin expression [72], suggesting that septins maintain mitochondrial stability in ciliates and free-living protists.

Measurements on cultured C2C12 cells suggested that Septin-7 is involved in myogenic cell division and myoblast fusion. This in turn indicated, based on previous experiments on satellite cells (see the review [73]), that it could play an important role in muscle regeneration, too. Indeed, we found (Figure 6C and 6E) that Septin-7 expression is increased following a mild muscle injury induced by the injection of BaCl_2_. The extent of regeneration following the injection was monitored by detecting the presence of Pax7 in satellite cells, and a clear correlation was seen in the time course of the expression pattern of the two molecules. In line with previous observations [74, 75] the most intense regeneration, and thus the increase in the expression of both Pax7 and Septin-7, was seen a few days after the injury, while after two weeks of regeneration was essentially complete and the expression of the two molecules returned to baseline. As the appearances of Pax7 as well as that of centrally located nuclei are accepted markers of muscle regeneration [76, 77], we concluded that the increased expression of Septin-7 is also an essential component of the regeneration process. Although these findings set the basis for the involvement of septin filaments in muscle repair further experiments are needed to elucidate their exact role in the process.

## Materials and methods

### Experimental model and subject details

#### Human Subjects

The human study was approved by the Ethics Committee of the Health Science Council, Budapest, Hungary (7917-1/2013/EKU 113/2013). Samples from *m. quadriceps femoris* of human patients going through amputation were taken and used in this study. Amputation surgery and biopsy preparation have been performed at the Kenézy Gyula Teaching Hospital of the University of Debrecen, Hungary.

#### Animal care and experimental design

Animal experiments were in compliance with the guidelines of the European Community (86/609/EEC). The experimental protocol was approved by the institutional Animal Care Committee of the University of Debrecen (2/2019/DEMAB). The mice were housed in plastic cages with mesh covers, and fed with pelleted mouse chow and water *ad libitum*. Room illumination was an automated cycle of 12 hours light and 12 hours dark, and room temperature was maintained within the range of 22–25°C.

#### Tamoxifen diet

Tamoxifen diet (*per os*) started immediately following separation from the mother at age of 3 weeks. Littermates were fed for 3 months without interruption. Chow (Envigo, TD 130857) contains 500 mg tamoxifen/kg diet, providing 80 mg tamoxifen/kg body weight per day assuming 20-25 g body weight and 3-4 g daily intake [78, 79].

#### Screening of HSA-MCM transgenic lines

B6.Cg-Tg(ACTA1-cre)79Jme/J strain originates from the Jackson Laboratory (Bar Harbor, Maine, USA). Mice hemizygous for this HSA-Cre79 transgene are viable, fertile, normal in size, and do not display any gross physical or behavioral abnormalities. These HSA-Cre79 transgenic mice have the *cre* recombinase gene driven by the human alpha-skeletal actin (*HSA* or *ACTA1*) promoter. Cre activity is restricted to adult striated muscle fibers. When bred with mice containing a *loxP*-flanked sequence of interest, Cre− mediated recombination will result in striated muscle-specific deletion of the flanked genome. Genomic DNA was isolated from finger biopsies [80] and screened for the presence of the HSA-MCM transgene by PCR using the appropriate primers spanned the C-terminus MerCre junction (see Table S1) and produced a 717-bp product.

#### Septin-7 genotyping and detection of exon4 deletion

C57BL/6J Septin-7^flox/flox^ (SS00) mice were obtained from Prof. Dr. Matthias Gaestel, Institute of Physiological Chemistry Hannover Medical School. These mice were crossed with B6.Cg-Tg(ACTA1-cre)79Jme/J mice (ssC0). The SsC0 offspring were back crossed with SS00 mice. Then SSC0 and SS00 mice were taken in breeding to produce litters for tamoxifen feeding. To investigate the effectivity of Septin-7 modification in the skeletal muscle of Cre+Septin-7flox/flox mice (Cre+ in the following), genomic DNA was prepared from *m. quadriceps femoris, m. biceps femoris* and *m. pectoralis*. Cre− Septin-7flox/flox mice did not have the MerCreMer construct, these mice were used as littermates. PCR primers were used to differentiate the flox from wild type mice, gene deletion was detected by the application of forward2 primer (see TableS1). Based on the presence of specific 151bp and 197bp bands, the wild type C57BL/6J (BL6 in the following) and floxed mice could be identified, while the deletion of exon4 proved to be partially achieved as represented by 256bp bands. According to the gene structure, exon4 (107bp, flox region is about 0.5kb) can be conditionally removed, which results in a non-sense mediated decay.

#### Cell culture

Mouse immortalized C2C12 myoblast cell line was cultured in High Glucose DMEM (Biosera, Nuaille, France) supplemented with 10% fetal bovine serum (Gibco by Life Technologies, Carlsbad, California, USA), 1% Penicillin-Streptomycin (Gibco) and 1% L-glutamine (Biosera). Medium was changed every other day and cells were subcultured at 80-90% confluence. Myoblast fusion and generation of differentiated myotubes was induced by exchanging the culture media to DMEM containing 2% horse serum (Gibco) approximately at 90% confluence.

#### CRISPR-Cas9

CRISPR/Cas9 knockout and HDR plasmid constructs specific to mouse Septin-7, targeting 3 different places in the coding sequence (in exon# 3, 4 and 5), were purchased from Santa Cruz BioTechnology. Transfection of the C2C12 cells with KO and HDR plasmids was carried out in serum-free Opti-MEM medium (Invitrogen by Thermo Fisher Scientific, Oregon, USA) using Lipofectamine 2000 transfection reagent (Invitrogen). 48 hours after transfection puromycin selection was applied for 5 days, then single cells were sorted according to expression of green fluorescent protein (GFP) and red fluorescent protein (RFP) from the KO and HDR vectors, respectively, using FACSAria flow cytometer (BD Biosciences, San Jose, CA, USA). Single cells were kept in normal culturing media until cell growth was observed in the appropriate cultures. Proliferation was continuously monitored by a transmission microscope.

#### Gene silencing

C2C12 cells were seeded in 6-well culture plates in DMEM containing 10% FBS. At 50-70% confluence, medium was replaced by serum-free OptiMEM and cells were transfected with Septin-7-specific shRNA constructs in retroviral pGFP-V-RS vectors (Origene, Cambridge, UK) using Lipofectamine 2000 transfection reagent. For controls, a non-effective scrambled shRNA cassette vector was employed (SCR). Three hours after transfection, the medium was replaced by complete DMEM, cells were allowed to recover and synthesize the coded shRNA for 48 hrs. Puromycin (2µg/ml) containing medium was applied to select cells containing the specific shRNA sequences until well defined cell clones were visible within the culture plates. Individual clones were then separated and cultured further to analyze the effective gene silencing by Western blot. Appropriate clones presenting significant Septin-7 expression change were tested throughout increasing passage numbers and continuously detectable lower Septin-7 expression was required to use the cell clone for further investigation (S7-KD cells).

## METHOD DETAILS

### *In vivo* experiments

#### *In vivo* CT imaging

For the in vivo imaging mice were anaesthetized by 3% isoflurane (Forane) with a dedicated small animal anaesthesia device. Whole body CT scans were acquired with the preclinical nanoScan SPECT/CT (Mediso Ltd, Hungary) scanner using the following acquisition parameters: X-ray tube voltage 60 kVp, current 86 mA; exposure time 170 ms per projection; voxel size: 1×1 mm. For the CT image reconstruction and analysis the Nucline and InterView™ FUSION software (Mediso LTD., Hungary) was used, respectively.

#### Voluntary activity wheel measurement

Mice from the different groups (see above) were housed in a cage with a mouse running wheel (Campden Instruments Ltd., Loughborough, UK). Wheels were interfaced to a computer and revolutions were recorded in 20 minute intervals, continuously for 14 days. The daily average and the maximal speed, the distance and the duration of running were calculated for each individual mouse and then averaged by groups (see Table S2).

#### Forepaw grip test

The force of forepaw was measured as described earlier [81]. Briefly, when the animals reliably grasped the bar of the grip test meter, they were then gently pulled away from the device. The maximal force before the animal released the bar was digitized at 2 kHz and stored by an online connected computer. The test was repeated 10-15 times to obtain a single data point. Measurements for the trained groups were always carried out before the 14 days running regime. For all other animal groups, the grip test was measured on the day when the animals were sacrificed.

### *In vitro* experiments

#### Measurement of muscle force

Muscle contractions were measured as described in our previous reports [82]. In brief, fast and slow twitch muscles, *m. extensor digitorum longus* (EDL) and *m. soleus* (Sol) were removed and placed horizontally in an experimental chamber continuously superfused (10 ml/min) with Krebs’ solution (containing in mM: NaCl 135, KCl 5, CaCl_2_ 2.5, MgSO_4_ 1, Hepes 10, glucose 10, NaHCO_3_ 10; pH 7.2; room temperature), equilibrated with 95% O_2_ plus 5% CO_2_. One end of the muscle was attached to a rod while the other to a capacitive mechano-electric force transducer (Experimetria, Budapest, Hungary). Two platinum electrodes placed adjacent to the muscle were used to deliver short, supramaximal pulses of 2 ms in duration to elicit single twitches. Force responses were digitized at 2 kHz using TL-1 DMA interface and stored with Axotape software (Axon Instruments, Foster City, CA, USA). Muscles were then stretched by adjusting the position of the transducer to a length that produced the maximal force response and allowed to equilibrate for 6 min.

Single pulses at 0.5 Hz were used to elicit single twitches. At least 10 twitches were measured under these conditions from every muscle. The individual force transients within such a train varied by less than 3% in amplitude, thus the mean of the amplitude of all transients was used to characterize the given muscle. To elicit a tetanus, single pulses were applied with a frequency of 200 Hz for 200 ms (EDL) or 100 Hz for 500 ms (Sol). Duration of individual twitches and tetani were determined by calculating the time between the onset of the transient and the relaxation to 10% of maximal force.

#### Isolation of single skeletal muscle fibers

Single muscle fibers from *m. flexor digitorum brevis* (FDB) were enzymatically dissociated in minimal essential media containing 0.2% Type I collagenase (Sigma) at 37°C for 30-40 min depending on the muscle weight [83, 84]. To release single fibers, the FDB muscles were triturated gently in normal Tyrode’s solution (1.8 mM CaCl_2_, 0 mM EGTA). The isolated fibers were then placed in culture dishes and stored at 4°C until use.

#### Immunofluorescent staining of isolated single fibers

To perform immunocytochemistry fibers were fixed immediately with 4% PFA for 20 minutes. After the fixation method 0.1 M glycine in PBS was used to neutralize excess formaldehyde. Fibers were permeabilized with 0.5% Triton-X (TritonX-100, Sigma) for 10 minutes and blocked with a serum-free Protein blocking solution (DAKO, Los Altos, CA, USA) for 30 minutes. Slides were rinsed three times with PBST solution. Primary antibodies (anti-RyR1, anti-Septin-7, and skeletal muscle-specific anti-α-blocking solution were added to the fibers and slides were incubated overnight at 4 °C in a humidity chamber. Samples were washed three times with PBST and incubated with fluorophore-conjugated secondary antibodies for 1 hour at room temperature. After three times washing a drop of mounting medium was added to each slide (Mowiol 4-88, Sigma) and coverslips placed on the mounting medium. Images from Alexa Fluor 488, TRITC, and DAPI labelled samples were acquired with an AiryScan 880 laser scanning confocal microscope (Zeiss, Oberkocken, Germany) equipped with a 20x air and a 40x oil objective. Excitation at 488 nm, 543 nm, and 405 nm wavelengths were used to detect fluorescence of the aforementioned secondary antibodies, respectively, while emission collected above 550 nm with a long pass filter.

#### Muscle regeneration

Skeletal muscle injury was accomplished by BaCl_2_ injection. 20 µl of 1.2% BaCl_2_ (dissolved in physiological saline) was injected to the left *m. tibialis anterior* muscle (TA) of BL6 mice (right *m. tibialis anterior* muscle was non-treated control). Mice were sacrificed 4 and 14 days later followed by removal of TA. Samples were obtained from both injected and non-injected muscles for Western blot analysis and for cryosection/paraffin embedded sections. Histological sections were stained with Hematoxylin and Eosin or DAB reaction was used to detect Pax7 expression.

#### Sample preparation for electron-microscopic studies

Freshly prepared TA was fixed *in situ* with fixative solution (3% glutaraldehyde in Millonig’s buffer). Small bundles of fixed muscle fibers were then postfixed in 1% OsO_4_ in water. For rapid dehydration of the specimens, graded ethanol followed by propylene-oxide intermediate was used. Samples were then embedded in Durcupan epoxy resin (Sigma). Ultrathin horizontal and transversal sections were cut using a Leica Ultracut UCT (Leica Microsystems, Wien, Austria) ultramicrotome and stained with uranyl acetate and lead citrate. Sections were examined with a JEM1010 transmission electron microscope (JEOL, Tokyo, Japan) equipped with an Olympus camera.

#### RT-PCR analysis

Cell cultures were dissolved, while human muscle biopsies and mouse skeletal muscle were homogenized in Trizol (Molecular Research Center, Cincinnati, OH, USA) with HT Mini homogenizer (OPS Diagnostics, Lebanon, NJ, USA) and subjected to a general RNA isolation protocol. In detail, after the addition of 20% chloroform samples were centrifuged at 4°C at 16,000×*g* for 15 min. Upper, aqueous phase of samples were incubated in 500 µL (cultures) or 750 µl (tissues) of RNase free isopropanol at room temperature for 10 min, then total RNA was harvested in RNase-free water and stored at –80°C. The assay mixture (20 μl) for reverse transcriptase reaction (Omniscript, Qiagen, Germantown, MD, USA) contained 1 µg RNA, 0.25 μl M), 1 μl High Affinity RT in 10 × RT buffer. Amplifications of specific cDNA sequences were carried out using specific primer pairs that were designed by Primer Premier 5.0 software (Premier Biosoft, Palo Alto, CA, USA) based on mouse and human nucleotide sequences published in GenBank and purchased from Bio Basic (Toronto, Canada). The specificity of custom-designed primer pairs was confirmed *in silico* by using the Primer-BLAST service of NCBI (http://www.ncbi.nlm.nih.gov/tools/primer-blast/). The sequences of primer pairs, the annealing temperatures for each specific primer pair, and the expected amplimer size for each polymerase chain reaction are shown in Table S1. Amplifications were performed in a programmable thermal cycler (Labnet MultiGene™ 96-well Gradient Thermal Cycler; Labnet International, Edison, NJ, USA) with the following settings: initial denaturation at 94°C for 1 min, followed by 30 cycles (denaturation at 94°C, 30 s; annealing at optimized temperatures for each primer pair for 30 s – see Table S1; extension at 72°C, for 60 s) and then final elongation at 72°C for 5 min. PCR products were mixed with EZ-Vision Dye 6X loading buffer and DNA bands were visualized following an electrophoresis in 1.2-2.5 % agarose gels.

#### Quantitative PCR analysis

Total RNA samples originating from different skeletal muscles (*m. tibialis anterior*, *m. pectoralis* and *m. quadriceps*) of mice were subjected to DNase treatment, and reverse transcription (Thermo Fisher Scientific, Waltham, MA, USA) according to the manufactureŕs instructions. Appropriate cDNA samples were used for quantitative PCR reaction (LightCycler 480, Roche, Basel, Switzerland) using either SYBRGreen mix (PCRBiosystems, Oxford, UK) and specific primer pairs for the detection of mitochondrial (16S RNA) and nuclear (hexokinase) RNA, or high-specificity Taqman assays (Mm00550197_m1) against mouse Septin-7 (Thermo Fisher Scientific).

#### Western blot analysis

Total cell lysates and skeletal muscle tissues were homogenized in a lysis buffer (20 mM Tris–HCl, 5 mM EGTA, Protease Inhibitor Cocktail (Sigma, Saint Louis, USA) with HT Mini homogenizer (OPS Diagnostics USA). Fivefold concentrated electrophoresis sample buffer (20 mM Tris–HCl, pH 7.4, 0.01% bromophenol blue dissolved in 10% SDS, 100 mM β-mercaptoethanol) was added to total lysates to adjust equal protein concentration of samples, and boiled for 5 min at 90°C. 8 or 10 μg of total protein (for Septin-7 and Pax7, respectively) were loaded to each lane, and separated in 7.5% SDS–polyacrylamide gel. Proteins were transferred to nitrocellulose membranes, blocked with 5% non-fat milk dissolved in phosphate saline buffer (PBS), then membranes were incubated with the appropriate primary antibodies overnight at 4°C (see Key resources table). After washing for 30 minutes in PBS supplemented with 1% Tween-20 (PBST), membranes were incubated with HRP-conjugated secondary antibodies (see Key resources table). Membranes were developed and signals were detected using enhanced chemiluminescence (Thermo Fisher Scientific). Optical density of signals was measured by ImageJ software (NIH, Bethesda, MD, USA) and results were normalized to the optical density of control tissues.

#### Immunofluorescent staining of cultured cells

To perform immunocytochemistry cell cultures were fixed immediately with 4% PFA for 15 minutes. After the fixation method 0.1 M glycine in PBS was used to neutralize excess formaldehyde. Fibers were permeabilized with 0.25% Triton-X (TritonX-100, Sigma) for 10 minutes and blocked with a serum-free Protein blocking solution (DAKO, Los Altos, CA, USA) for 30 minutes. Slides were rinsed three times with PBST solution. Primary Septin-7 antibody was diluted in blocking solution, added to the samples, and slides were incubated overnight at 4 °C in a humidity chamber. On the next day samples were washed three times with PBST and incubated with fluorophore-conjugated secondary antibodies and TRITC-phalloidin (1:1000) for 1 hour at room temperature. After three times washing a drop of mounting medium was added to each slide. Confocal images were acquired with an AiryScan laser scanning confocal microscope as described above.

#### Determination of cellular proliferation

The degree of cellular growth (reflecting proliferation) was determined by CyQUANT NF Cell Proliferation Assay Kit (Invitrogen). C2C12 cells (2500 cells/well) were cultured in 96-well black plates with clear bottoms (Greiner Bio-One, Mosonmagyaróvár, Hungary) for 48 hours. HBSS buffer was prepared (Component C) with deionized water, then one time diluted (1x) dye binding solution was added to the CyQUANT NF dye reagent (Component A). Growth medium was exchanged to 100 μL of 1x dye binding solution. Microplate was covered and incubated at 37°C for 30 minutes, and fluorescence was measured at 485 nm excitation and 530 nm emission wavelengths using FlexStation 3 multimode microplate reader (Molecular Devices, San Jose, CA, USA). Relative fluorescence values were expressed as percentage of control regarded as 100%.

#### Fusion index

Progress of myotube differentiation was quantified using immunocytochemistry in C2C12 cells. Cells were plated into glass coverslips, in each day of differentiation process samples were fixed in 4% paraformaldehyde (PFA) solution, and subjected to a desmin-specific immunolabeling and DAPI staining. Confocal images were taken from the appropriate samples. Fusion index was calculated as the ratio of the nuclei number in myocytes with two or more nuclei versus the total number of nuclei within the visual fields.

### Quantification and statistical analysis

#### Statistical analysis

Pooled data were expressed as mean±standard error of the mean (SEM). The differences between control mice and animals on tamoxifen diet were assessed using one-way analysis of variance (ANOVA) and all pairwise Bonferroni’s multiple comparison method using the statistical program Prism (GraphPad Software, San Diego, CA, USA). T-test was used to test the significance and a p value of less than 0.05 was considered statistically significant.

## Supporting information

Supplemental informations

## Acknowledgements

We are grateful to Matthias Gaestel (Institute of Physiological Chemistry, Hannover Medical School, Hannover, Germany) for providing the loxP mice for the experiments. The authors thank Mrs. R. Őri for her excellent technical assistance. Authors wish to thank László Balkai, Ildikó Garai and Scanomed Ltd. (Scanomed Translational Centre, Debrecen, Hungary) for *in vivo* imaging.

