## Supplemental informations for "Septin-7 is indispensable for proper skeletal muscle architecture and function"

**This PDF file includes:**

Key resources Table
Tables S1 to S2
Figures S1 to S8 with appropriate legends

**Supplemental figures and tables**

**KEY RESOURCES TABLE**

| REAGENT or RESOURCE | SOURCE | IDENTIFIER |
|---|---|---|
| Antibodies | | |
| rabbit monoclonal anti-Septin-7 | IBL | Cat#JP18991 RRID:AB_1630825 |
| mouse monoclonal anti-α-actinin | Santa Cruz BioTechnology | Cat#sc-17829 RRID:AB_626633 |
| mouse monoclonal anti-Pax7 | Sigma | Cat#SAB1404168 RRID:AB_10738723 |
| rabbit polyclonal anti-desmin | Sigma | Cat#D8281 RRID:AB_476910 |
| mouse monoclonal anti-Ryanodin receptor 1 | Thermo Fisher Scientific | Cat#MA3-925 RRID:AB_2254138 |
| mouse monoclonal anti-α-actinin (skeletal muscle specific) | Sigma | Cat#A7811 RRID:AB_476766 |
| goat monoclonal HRP-conjugated anti-rabbit | Bio-Rad Laboratories | Cat#170-6515 RRID:AB_11125142 |
| goat monoclonal HRP-conjugated anti-mouse | Bio-Rad Laboratories | Cat# 170-6516 RRID:AB_11125547 |
| goat monoclonal Alexa Fluor 488 conjugated anti-rabbit IgG | Thermo Fisher Scientific | Cat# A32731 RRID:AB_2633280 |
| goat monoclonal Cyanine3 conjugated anti-Mouse IgG | Thermo Fisher Scientific | Cat# A10521 RRID:AB_1500665 |
| Bacterial and virus strains | | |
| JM 109 competent *E. coli* cells | Promega | Cat# L2005 |
| Biological samples | | |
| human *m. quadriceps femoris* | Kenézy Gyula Teaching Hospital of the University of Debrecen | https://kenezykorhaz.unideb.hu/en |
| Chemicals, peptides, and recombinant proteins | | |
| Tamoxifen containing chow | Envigo | Cat# TD 130857 |
| Type I collagenase | Sigma | Cat#SCR103 |
| Isoflurane | Forane | Cat# NDC 10019-360 |
| Protein blocking solution serum-free | Dako | Cat#X0909 |
| TRITC-phalloidin | Sigma | Cat#P1951 |
| Mowiol 4-88 | Sigma | Cat# **81381** |
| Durcupan epoxy resin | Sigma | Cat#44611 |
| EZ-Vision Dye 6X | VWR Life Science | Cat# 97064-190 |
| Lipofectamine 2000 | Thermo Fisher Scientific | Cat#11668019 |
| Opti-MEM Reduced Serum medium | Thermo Fisher Scientific | Cat# 31985070 |
| Critical commercial assays | | |
| Omniscript RT kit | Qiagen | Cat# 205113 |
| RNase free DNase kit | Thermo Fisher Scientific | Cat# EN0521 |
| SYBRGreen mix | PCRBiosystems | Cat#PB20.11-51 |
| High-specificity Taqman assays | Thermo Fisher Scientific | Cat#4331182 |

| | | |
|---|---|---|
| CyQUANT NF Cell Proliferation Assay Kit | Invitrogen | Cat# C35006 |
| DAB substrate kit | Thermo Fisher Scientific | Cat#34002 |
| SuperSignal™ West Pico PLUS Chemiluminescent Substrate | Thermo Fisher Scientific | Cat# 34577 |
| Deposited data | | |
| Human reference genome NCBI | Genome Reference Consortium | http://www.ncbi.nlm.nih.gov/projects/genome/assembly/human |
| Mouse reference genome NCBI | Genome Reference Consortium | http://www.ncbi.nlm.nih.gov/projects/genome/assembly/mouse |
| Experimental models: Cell lines | | |
| Mouse immortalized C2C12 | ATCC | Cat# CRL-1772 RRID:CVCL_0188 |
| Experimental models: Organisms/strains | | |
| B6.Cg-Tg(ACTA1-cre)79Jme/J mice | Jackson Laboratory | Cat#006149 RRID:IMSR_JAX:006149 |
| C57BL6 Septin-7$^{flox/flox}$ (SS00) mice | Prof. Dr.Matthias Gaestel | Institute of Physiological Chemistry Hannover Medical School |
| Oligonucleotides (see Table S1.) | | |
| Recombinant DNA | | |
| Septin-7 specific CRISPR/Cas9 KO plasmid constructs | Santa Cruz BioTechnology | Cat#sc-433427 |
| Septin-7 specific HDR plasmid constructs | Santa Cruz BioTechnology | Cat#sc-433427-HDR |
| Septin-7-specific shRNA constructs in retroviral pGFP-V-RS vectors | Origene | Cat#TR30007 |
| Software and algorithms | | |
| Nucline and InterView™ FUSION software | Mediso Ltd. | http://ctamed.com/en/portfolio/software-solutions/#fusion |
| Axotape software | Axon Instruments | https://www.bioz.com/result/axotape/product/Molecular%20Devices%20Llc |
| Primer Premier 5.0 software | Premier Biosoft | http://downloads.fyxm.net/Primer-Premier-101178.html |
| Primer-BLAST | | https://www.ncbi.nlm.nih.gov/tools/primer-blast/ |
| Statistical program Prism | GraphPad Software | https://www.graphpad.com/scientific-software/prism/ |
| Image J1.40g | Schneider et al., 2012 | https://imagej.nih.gov/ij/download.html |

| | | |
|---|---|---|
| Origin 8.6 | Originpro | https://originpro.informer.com/8.6/ |
| Experimental devices | | |
| FACSAria flow cytometer | BD Biosciences | https://www.bdbiosciences.com/en-eu/instruments/research-instruments/research-cell-sorters/facsaria-iii |
| nanoScan SPECT/CT | Medisol Ltd | http://scanomedtranslational.com/ |
| Mouse running wheel | Campden Instruments Ltd | Bodnar et al, 2014 |
| Capacitive mechano-electric force transducer | Experimetria | Bodnar et al, 2016 |
| Leica Ultracut UCT | Leica Microsystems | https://www.leica-microsystems.com |
| JEM1010 transmission electron microscope | JEOL | https://www.jeolusa.com/PRODUCTS/Transmission-Electron-Microscopes-TEM |
| HT Mini homogenizer | OPS Diagnostics | Cat#BM-D1030E |
| Labnet MultiGene™ 96-well Gradient Thermal Cycler | Labnet International | Cat# TC9610-230 |
| LightCycler 480 | Roche | Cat# 05015243001 |
| FlexStation 3 multimode microplate reader | Molecular Devices | Cat#Flex3 |
| AiryScan Confocal microscope | Zeiss | https://www.zeiss.com/microscopy/int/dynamic-content/news/2014/news-lsm-880.html |

**Table S1.** Nucleotide sequences, amplification sites, GenBank accession numbers, amplimer sizes and PCR reaction conditions for each primer pair are shown. *Related to Figure 1; Figure 2; and Figure S1.*

| *Gene* | *Primer* | *Nucleotide sequence (5'→3')* | *GenBank Accession No.* | *Annealing temperature* | *Amplimer size (bp)* |
|---|---|---|---|---|---|
| **SEPT1** | sense | GGGTTTGACTTCACGCTAATG (101-121) | **NM_001365977** | 58 °C | 443 |
| **Homo sapiens** | antisense | CCAATGACTGGGATGATGTTG (523-543) | | | |
| **SEPT2** | sense | GCGAAGATTCTCATTACC (367–384) | **NM_001008491** | 52 °C | 476 |
| **Homo sapiens** | antisense | TACCACTGTCAGGCGTAG (825–842) | | | |
| **SEPT3** | sense | TGAATGTTTGCGAATGTTG (4315-4333) | **NM_019106** | 55 °C | 220 |
| **Homo sapiens** | antisense | GATTGGCTGGGACTGGTA (4517-4534) | | | |
| **SEPT4** | sense | CCGAAAGTCCGTGAAGAA (698-715) | **NM_004574** | 55 °C | 221 |
| **Homo sapiens** | antisense | TGTCCACAATGGTGAGCC (901-918) | | | |
| **SEPT5** | sense | CTCAACCGAAAGAACATCCAA (513-533) | **NM_002688** | 55°C | 319 |
| **Homo sapiens** | antisense | TGCCTATAACGGCGAAGG (814-831) | | | |
| **SEPT6** | sense | TTCACGCTTAAACAACCA (3403-3420) | **NM_145799** | 52°C | 407 |
| **Homo sapiens** | antisense | CCTCGTATCCAGGAATGTA (3791-3809) | | | |
| **SEPT7** | sense | CTTATTGCCAAAGCAGAC (798-815) | **NM_001788** | 50°C | 434 |
| **Homo sapiens** | antisense | AGAGGGCTCTTAGTCAGC (1214-1231) | | | |
| **SEPT8** | sense | TGCCTCTACTTCATCACGC (559-577) | **NM_001098811** | 56 °C | 493 |
| **Homo sapiens** | antisense | TGTCACCATCGCTGTCCT (1034-1051) | | | |
| **SEPT9** | sense | AGGGCACAGATGACCAAAG (3043-3061) | **NM_001113495.1** | 56°C | 287 |
| **Homo sapiens** | antisense | GGCAAGTCGGCAAAGTAAA (3311-3329) | | | |
| **SEPT10** | sense | CTGGCAACAGGCAGCAAC (1449-1466) | **NM_001321509.2** | 56°C | 366 |
| **Homo sapiens** | antisense | AGCAAAGGTGTCGGGTGG (1797-1814) | | | |
| **SEPT11** | sense | AGGATGATGGGCTTTCTA (2449-2466) | **NM_001306147** | 50 °C | 279 |
| **Homo sapiens** | antisense | CTTACGGGTCATGTTGTG (2710-2727) | | | |
| **SEPT12** | sense | ATGTGGTGCCCGTGATTG (513–530) | **NM_001154458** | 56 °C | 324 |
| **Homo sapiens** | antisense | GGAGGTGGGAGCGGATAA (819–836) | | | |
| **SEPT14** | sense | GAAGAAACTGCTGCTCAA (786-803) | **XM_011515373** | 50 °C | 488 |
| **Homo sapiens** | antisense | TTCCTTATCTCCTCCTGT (1256-1273) | | | |
| **SEPT1** | sense | CACGGCACAAACTCTGACC (222-240) | **NM_017461** | 56°C | 500 |
| **Mus musculus** | antisense | CAACGACTGCGAAAGGGA (704-721) | | | |
| **SEPT2** | sense | GAACAGGCGTCACATCA (550-566) | **NM_001159719** | 50 °C | 352 |
| **Mus musculus** | antisense | AACCTTCTTGCCTTTGG (885-901) | | | |
| **SEPT3** | sense | CCACTGCTGCCTCTACT (776-792) | **NM_001358836** | 52 °C | 231 |
| **Mus musculus** | antisense | TTGTCCTCCAAATCCTC (990-1006) | | | |
| **SEPT4** | sense | GCAAACCGTGGAGATTA (654-670) | **NM_011129** | 52 °C | 325 |
| **Mus musculus** | antisense | CGCCTTAGCCAAGATAG (962-978) | | | |
| **SEPT5** | sense | CGCAAGTCCGTCAAGAAA (194-211) | **NM_213614** | 55 °C | 268 |

| | | | | | |
|---|---|---|---|---|---|
| **Mus musculus** | antisense | TGGGCTTCCAACATTCAG (444-461) | | | |
| **SEPT6** | sense | ACATCATTCCCGTTATTGC (820-838) | **NM_001177324** | 52°C | 221 |
| **Mus musculus** | antisense | CATCATCTTGTTGCCTATCTT (1020-1040) | | | |
| **SEPT7** | sense | CCTTGAGGGCTATGTGGG (278-295) | **NM_009859** | 56°C | 250 |
| **Mus musculus** | antisense | CAGCAGCAACTGAACACCAC (508-527) | | | |
| **SEPT8** | sense | CACAGAGGAGGTGAAGGT (1000-1017) | **NM_033144** | 54°C | 471 |
| **Mus musculus** | antisense | TTGAAGGCGTTGGTCTC (1454-1470) | | | |
| **SEPT9** | sense | AACATTGTCCCAGTCATCG (1400-1418) | **NM_001113486** | 54°C | 381 |
| **Mus musculus** | antisense | GCGTTTCACTCGGTAGG (1764-1780) | | | |
| **SEPT10** | sense | GACACGACCTCCAAGAT (1088-1104) | **NM_001024910** | 52°C | 349 |
| **Mus musculus** | antisense | GACGCTCACCATAGAAC (1420-1436) | | | |
| **SEPT11** | sense | CCTGTGCGTGGGTGAGA (287-303) | **NM_001310669** | 55 °C | 433 |
| **Mus musculus** | antisense | GATGGTGTCGGCTTTCG (703-719) | | | |
| **SEPT12** | sense | AGTCCAAAGTATGGCAGTC (344-362) | **NM_027669** | 52°C | 105 |
| **Mus musculus** | antisense | GCTTCAATCCCTTCTCC (432-448) | | | |
| **SEPT14** | sense | TCCACTCTTGGGCATTT (160-176) | **NM_028826** | 50 °C | 290 |
| **Mus musculus** | antisense | CTGGCTTCCTTGTTTATCT (431-449) | | | |
| **HSA-MCM** | sense | GCATGGTGGAGATCTTTGA | | 58 °C | 717 |
| **Mus musculus** | antisense | CGACCGGCAAACGGACAGAAGC | | | |
| **SEPT7** | sense1 | CTTTGCACATATGACTAAGC | | 58 °C | 151 WT |
| **WT-loxP-KD** | sense2 | GCTTCTTTTATGTAATCCAGG | | | 197 Flox |
| **Mus musculus** | antisense | GGTATAGGGGACTTTGGGG | | | 256 KD |
| **GAPDH** | sense | AAGGTCGGAGTCAACGGATTTGG(99-121) | **NM_001289726.1** | 58 °C | 322 |
| **Mus musculus** | antisense | AATGAGCCCCAGCCTTCTCCAT (399-420) | | | |

**Table S2: Voluntary running parameters** *Related to Figure 2; and Figure S3.*

3 month Tamoxifen treatment 4 week after birth (4 month old)

10 days long running experiment **during Tamoxifen treatment**

| Parameters of running | BL6 n=3 | Cre- n=2 | Cre+ n=4 |
|---|---|---|---|
| **Distance (km)** | 2.46±0.19 | 2.34±0.06 | 0.83±0.02*** |
| **Duration (min)** | 343.3±15.2 | 317.1±11.1 | 264.4±4.2*** |
| **Average speed (m/min)** | 7.27±0.44 | 7.30±0.32 | 4.01±0.13*** |
| **Maximal speed (m/min)** | 14.04±0.56 | 15.71±0.62 | 10.31±0.20*** |

*** shows significant difference from control at $p<0.001$.

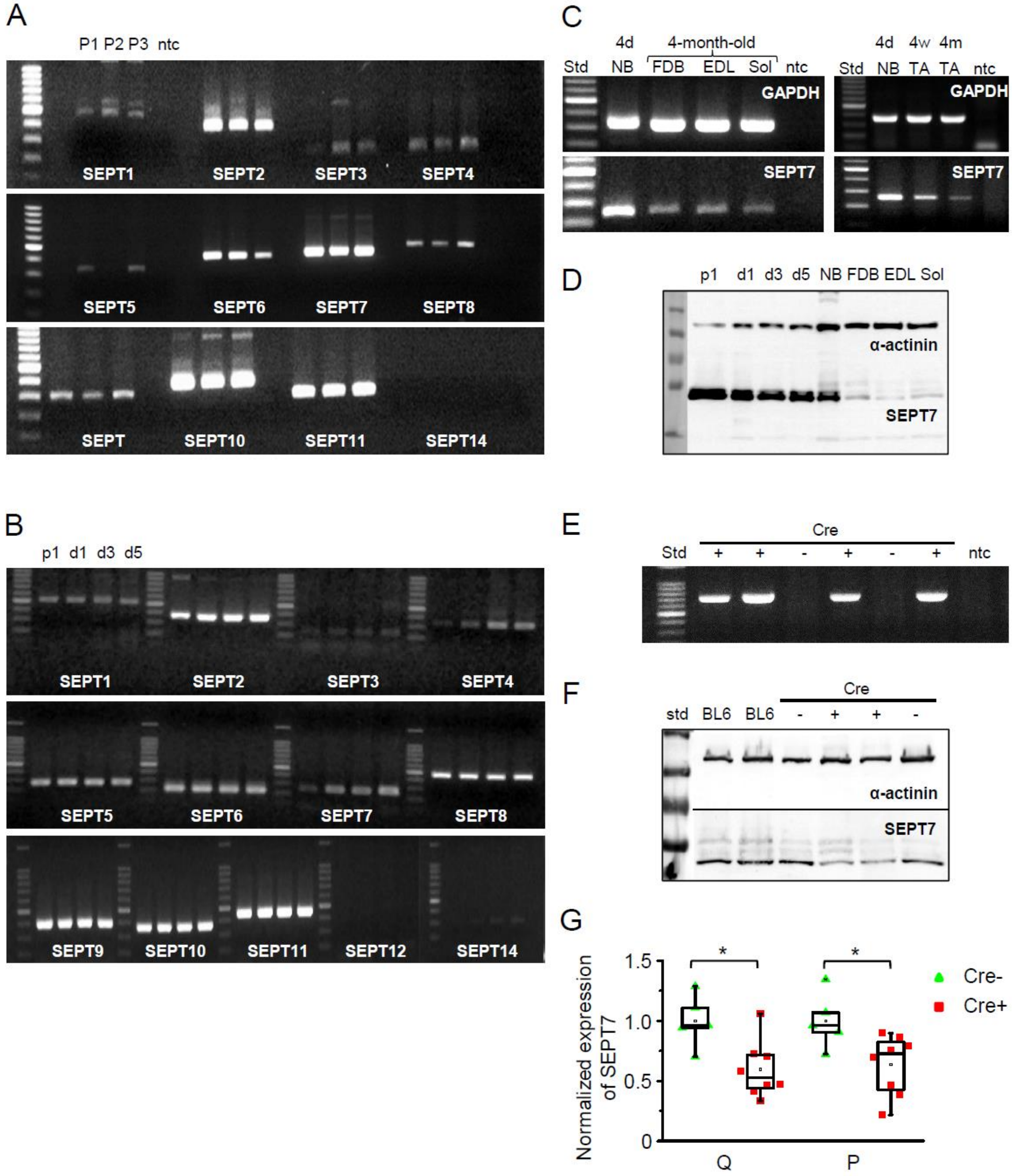

**Figure S1.** (**A) Expression of different septin isoforms at mRNA level in human skeletal muscle (*m. quadriceps femoris*). Related to Figure 1 and Figure 2.** Independent samples from 3 patients (P1-P3) were examined. Extended experiment of Fig 1A**. (B)** mRNA expression of all septin isoforms in differentiating C2C12 myoblast cells. Proliferating cells (p1) and myotubes at different stages of development (d1, d3, and d5) are demonstrated**. (C)** Age related (age of 4 days-

4d, 4 weeks-4w, 4 month-4m) mRNA expression of SEPTIN7 in different skeletal muscle types of mouse. GAPDH was used as loading control; NB: newborn muscle, TA: *m. tibialis anterior,* FDB: *m. flexor digitorum brevis,* EDL: *m. extensor digitorum longus* and *Sol: m. soleus* were examined**.** **(D)** Ontogenesis dependent Septin-7 protein expression in differentiating C2C12 cells, newborn and different muscle types of adult (4 months old) mice, α-actinin was used as control. (**E**) Screening for the presence of the HSA-MCM transgene from genomic DNA by PCR. (**F**) Altered Septin-7 expression in Cre+ mice following a tamoxifen diet as compared to Cre- and control BL6 mice. The *m. quadriceps femoris* (Q) and *m. pectoralis* (P) were analyzed; α-actinin was used as a normalizing control. (**G**) Pooled data of the Septin-7 protein expression in different muscle types of tamoxifen fed mice. 8 Cre+ and 5 Cre- were examined from 3 independent litters. Data represent mean ± standard error of the mean, $*p < 0.05$.

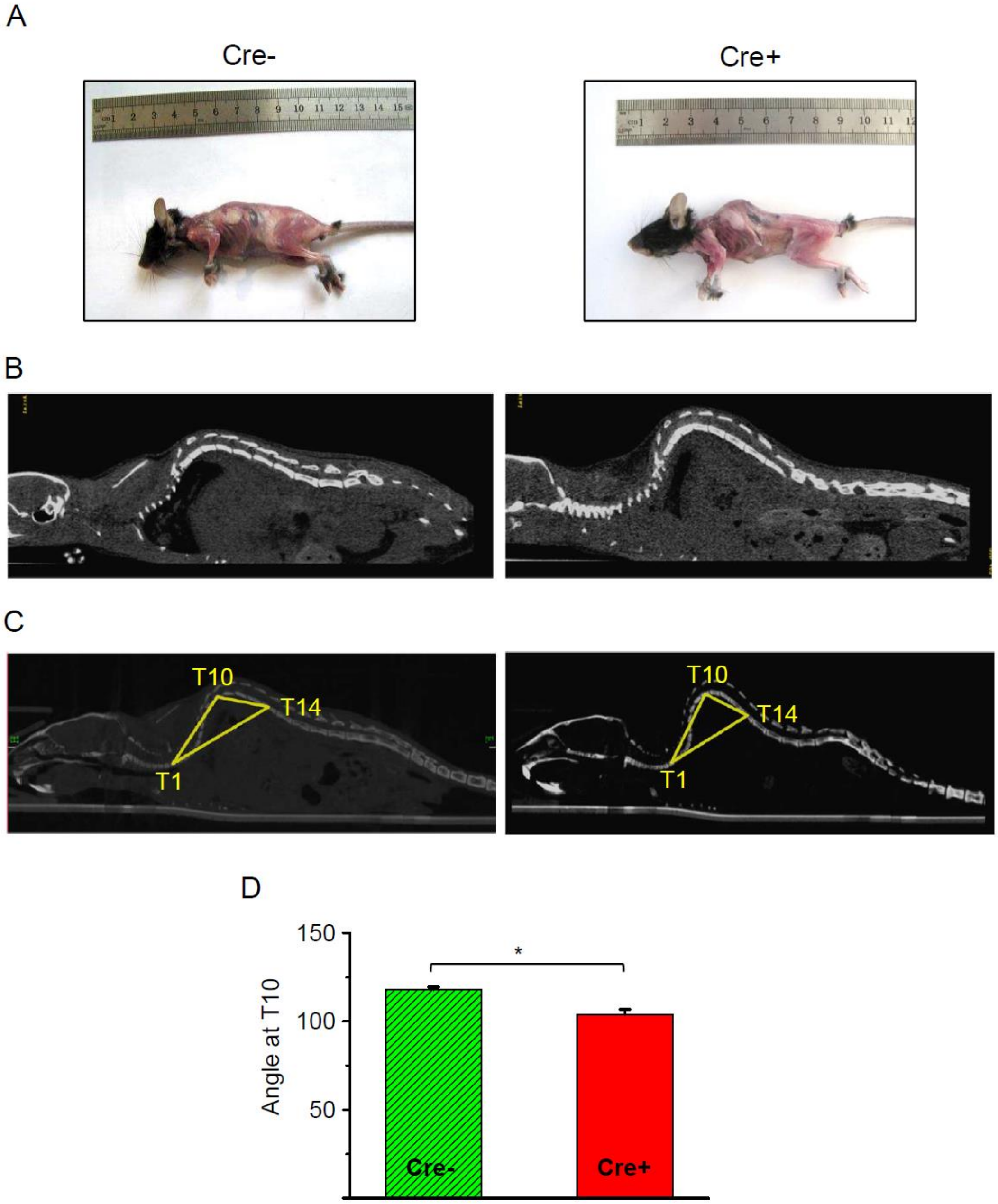

**Figure S2. Effects of Septin-7 knock-down on the phenotype of Cre+ mice.** (**A**) **Related to Figure 2.** Images of a tamoxifen-fed control (Cre-) and Cre+ mice. (**B**) Coronal CT image of a Cre- and Cre+ mouse. (**C**) Reconstruction of the whole backbone from CT images. A triangle was drawn to the thoracic 1$^{st}$ (T1), 10$^{th}$ (T10), and 14$^{th}$ (T14) vertebra and the angle at T10 was determined. (**D**) Average of angles at T10 from 3 Cre- and 3 Cre+ mice, *p<0.05.

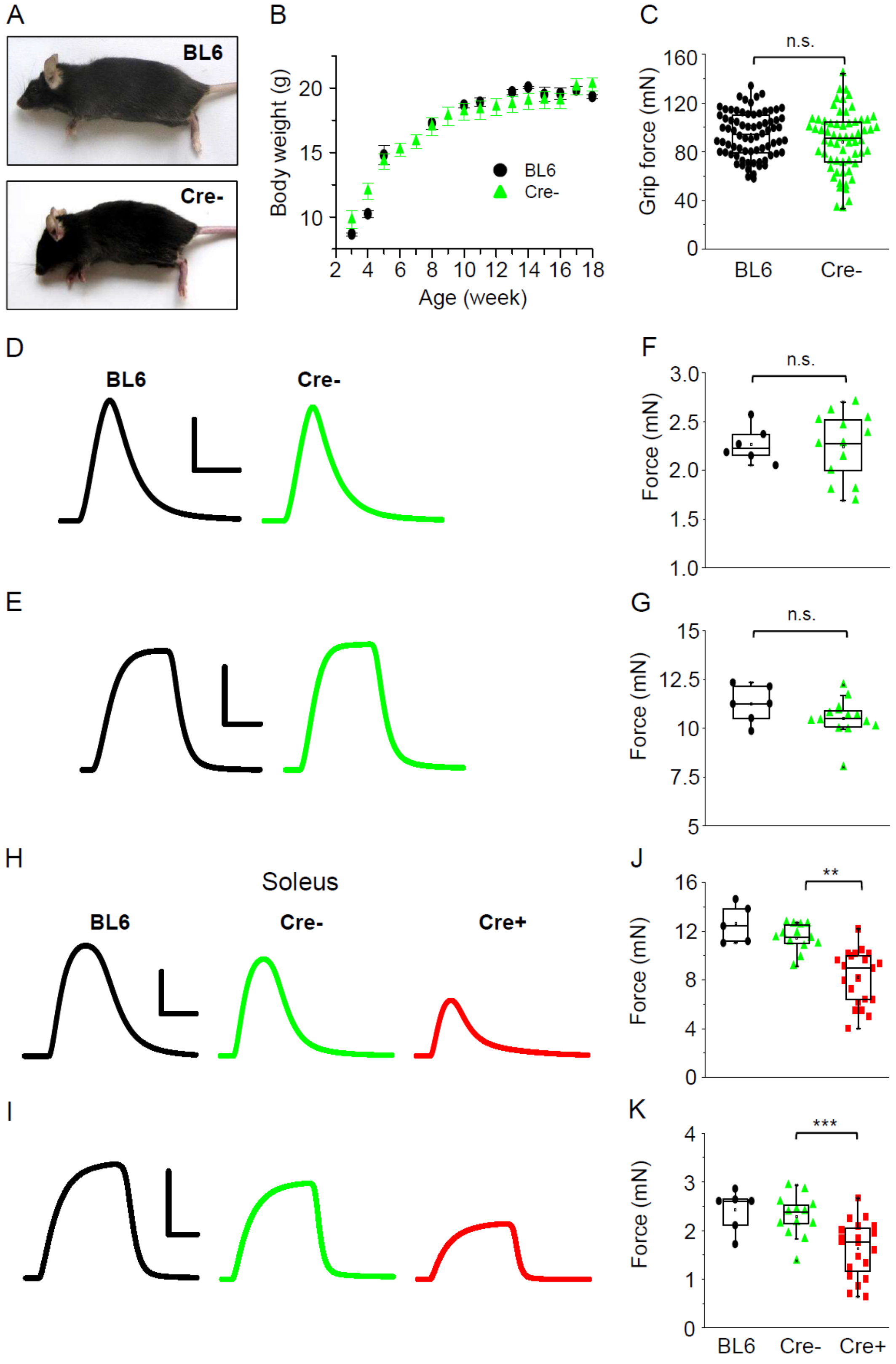
A
BL6
Cre-
B
Body weight (g)
20
15
10
BL6
Cre-
2 4 6 8 10 12 14 16 18
Age (week)
C
n.s.
160
120
80
40
0
Grip force (mN)
BL6
Cre-
D
BL6
Cre-
F
n.s.
3.0
2.5
2.0
1.5
1.0
Force (mN)
E
G
n.s.
15
12.5
10
7.5
5
Force (mN)
H
Soleus
BL6
Cre-
Cre+
J
**
16
12
8
4
0
Force (mN)
I
K
***
4
3
2
1
0
Force (mN)
BL6
Cre-
Cre+

**Figure S3. Effects of Tamoxifen feeding on the phenotype of BL6 and Cre- mice.** (**A**) **Related to Figure 2.** Images of tamoxifen-fed BL6 and Cre- mice. (**B**) Body weight increase in BL6 (black circle, n=3) and Cre- (green triangle, n=11) mice. (**C**) Grip force in BL6 (n=3) and Cre- (n=4) mice. Representative twitch **(D)** and tetanic force **(E)** transients in EDL. Peak twitch **(F)** and tetanic force **(G)** in EDL from BL6 (n=3) and Cre- (n=7) mice. Representative twitch **(H)** and tetanic force **(I)** transients in *m. soleus* (Sol). Peak twitch **(J)** and tetanic force **(K)** in Sol from BL6 (n=3), Cre- (n=7) and Cre+ (n=11) mice. Calibration in panel D: 1 mN and 50 ms; E: 5 mN and 100 ms; H: 1 mN and 100 ms; I: 10 mN and 200 ms, ** $p<0.01$, *** $p<0.001$.

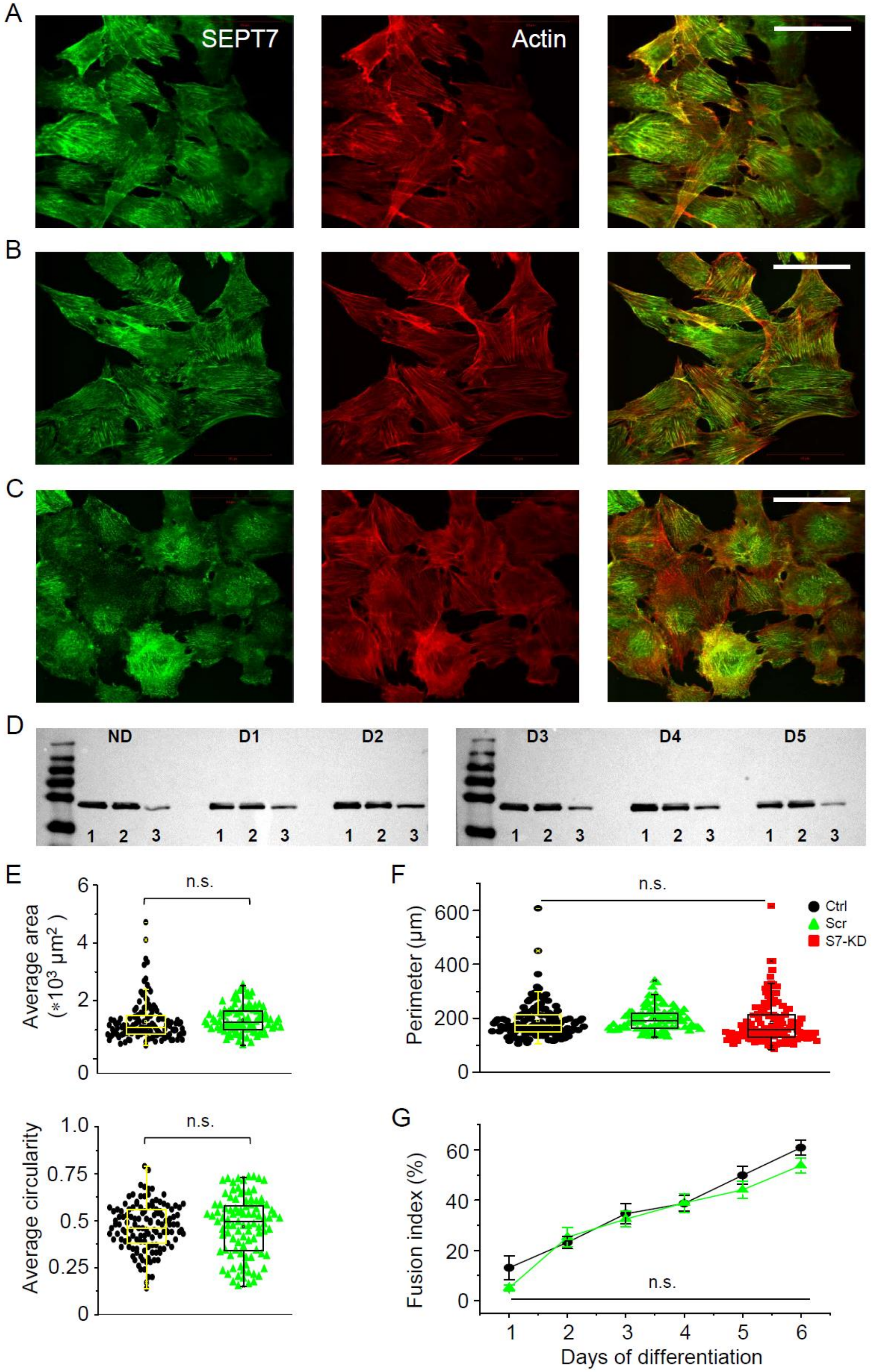
A
SEPT7
Actin
B
C
D
ND
D1
D2
D3
D4
D5
1 2 3
E
n.s.
Average area (*10³ µm²)
Average circularity
F
n.s.
Perimeter (µm)
Ctrl
Scr
S7-KD
G
Fusion index (%)
n.s.
Days of differentiation

**Figure S4. Septin-7 filament structure and the effect of gene silencing on cellular parameters. Related to Figure 3.** Intracellular localization of Septin-7 (green) and actin (red) filaments individually and merged in control (Ctrl, **A**), scrambled (Scr, **B**), and S7-KD (**C**) C2C12 cells. Calibration is 50 μm. (**D**) Modified expression of Septin-7 protein (50kDa) was followed during the differentiation program of Ctrl, Scr and S7-KD cells. ND refers to non-differentiated, D1-D5 to appropriate days (1-5) following the induction of differentiation, while numbers 1, 2, and 3 at the bottom of the gels indicate Ctrl, Scr, and S7-KD samples, respectively. (**E**) Quantitation of change in S7-KD cell morphology (area and circularity). Black circles represent data from control, while green triangles from scrambled cells. Individual data points (symbols) and average values (boxes with error bars) are shown for each group. The number of cells investigated was 127 in Ctrl, 96 in Scr, and 121 in S7-KD cultures; the experiment was repeated twice (N=2). (**F**) Average perimeter was also calculated from different cells and this data did not show significant difference between Ctrl (black), Scr (green) and S7-KD samples (red). (**G**) Differentiation during 6 days of culturing was assessed by calculating the fusion index in Ctrl (black) and Scr (green) cultures revealed to be not significantly different (n=20; N=2).

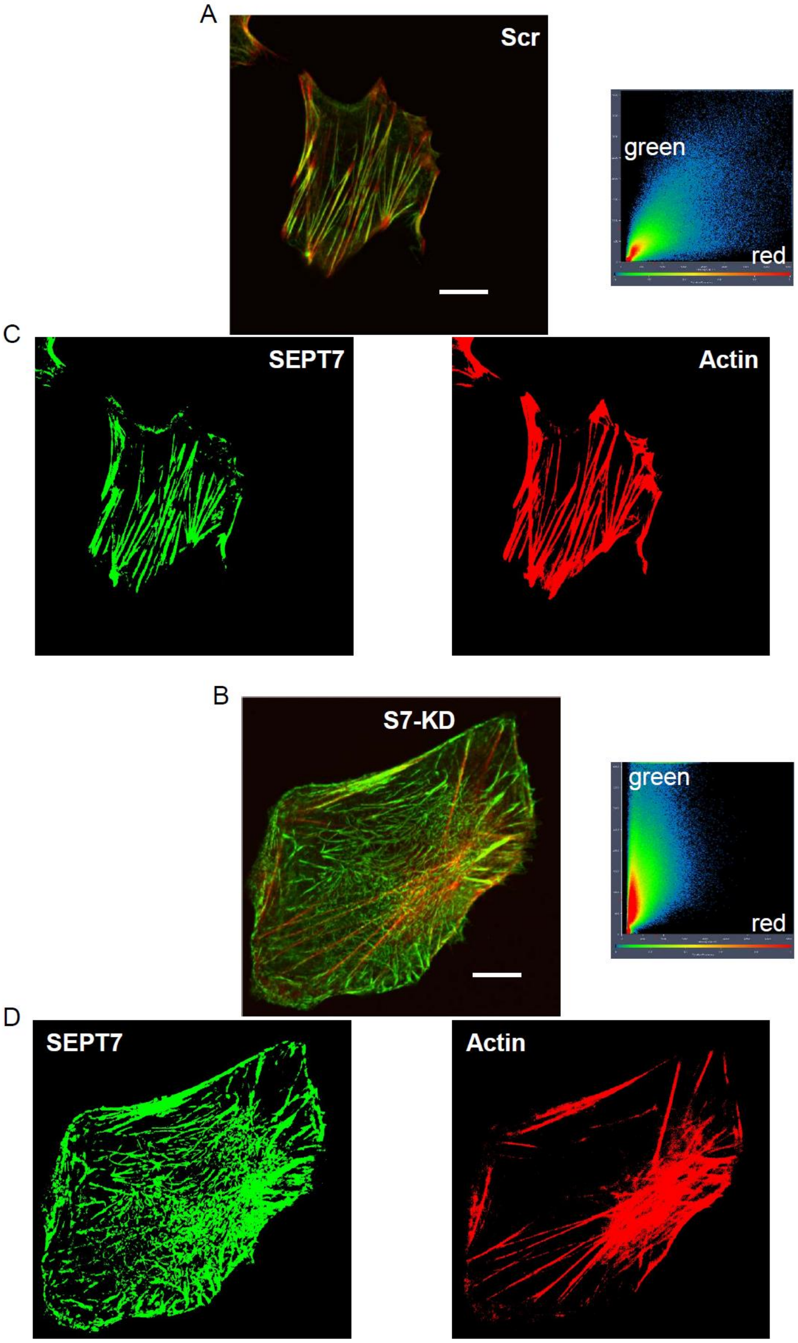

**Figure S5. SEPT 7 downregulation modifies the intracellular architecture of C2C12 cells. Related to Figure 3.** Representative confocal images of Septin-7 (green) and actin (red) filaments in Scr (**A**) and a S7-KD (**B)** C2C12 cell, where colocalization analysis was also performed. Note the asymmetric distribution of intensities in case of S7-KD cells. Filamentous images (**C, D**) were created with thresholding of the separate channels. Scale bar in panel A and B corresponds to 10 μm.

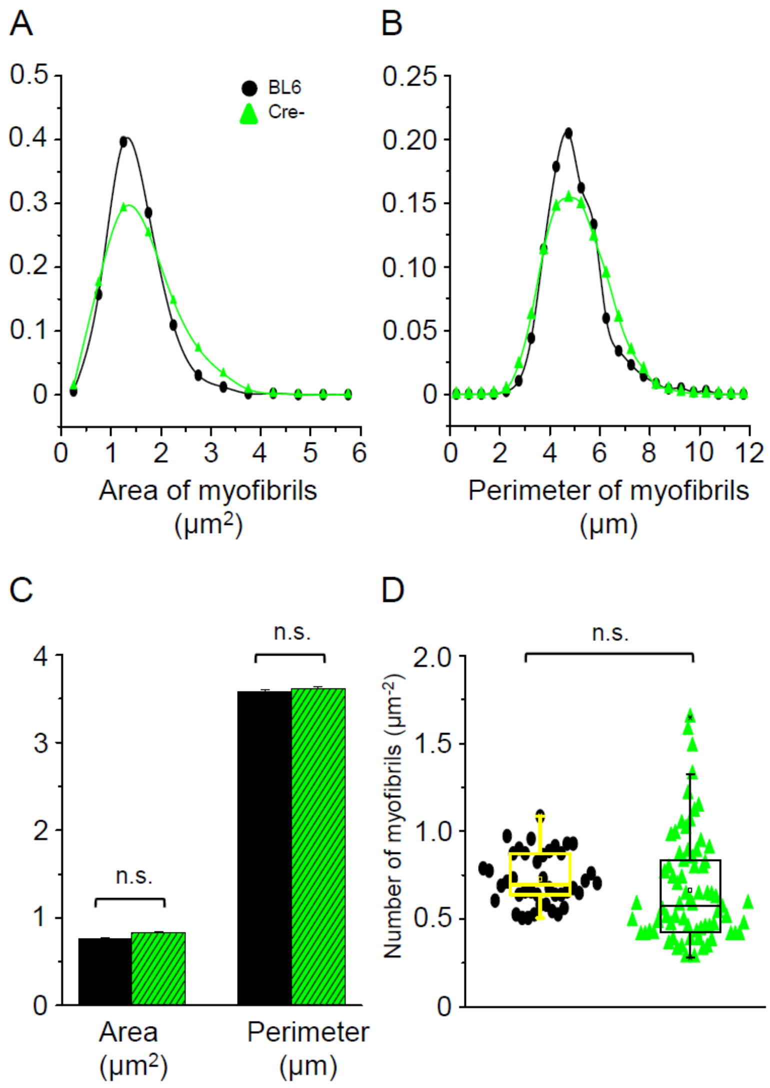

**Figure S6. Changes in myofibrillar parameters with *in vivo* knockdown of Septin-7. Related to Figure 4.** Myofibrillar parameters calculated from cross sectional EM pictures of TA muscles of control BL6 (black), and Cre- (green) mice. Histograms represent the distribution of mitochondrial area (**A**), and perimeter (**B**) in control, and Cre- samples, respectively. (**C**) Average of mitochondrial area and perimeter. Number of individual myofibrils (for area and perimeter) was 1916 and 3012, respectively (**D**). The number of myofibrils within 1 μm$^2$ of visual field in control BL6 and Cre- mice. The number of visual fields used for the calculation of myofibrillar data was 42 and 72 in samples originated from BL6 and Cre- mice, respectively.

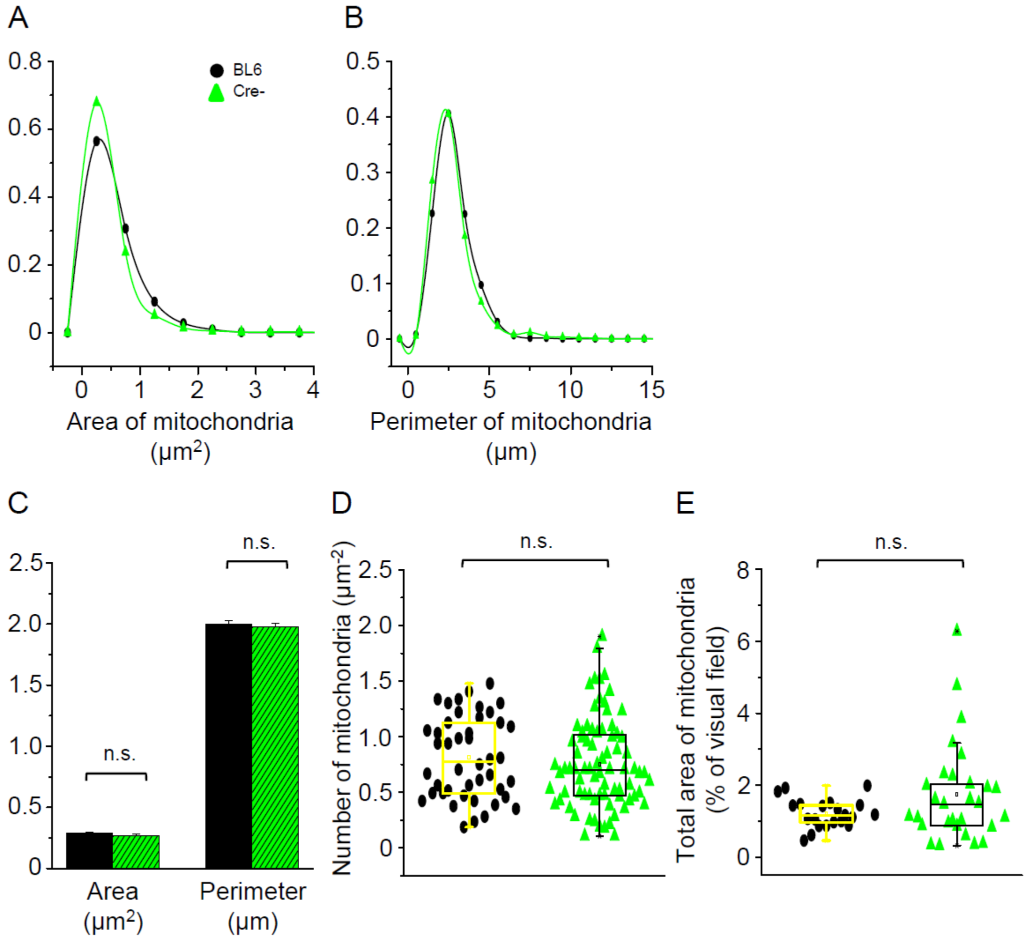

**Figure S7. Changes in mitochondrial parameters with *in vivo* knockdown of Septin-7. Related to Figure 5.** Mitochondrial parameters calculated from cross or longitudinal EM sections of TA muscles from control (black), and Cre- (green) mice. Distribution of mitochondrial area (**A**) and perimeter (**B**) calculated from the transversal muscle sections of control BL6 and Cre- mice, the average of these parameters is presented in panel **C**. The numbers of individual mitochondria (for area and perimeter) were 877 and 1589, respectively. **(D)** The number of mitochondria within 1 μm$^2$ of visual field in samples from BL6 control and Cre- mice. The number of visual fields for the calculation were 44 and 81, respectively. **(E)** The total area of mitochondria within the actual visual fields (n=21 and 29, respectively) in samples of control and Cre- animals.

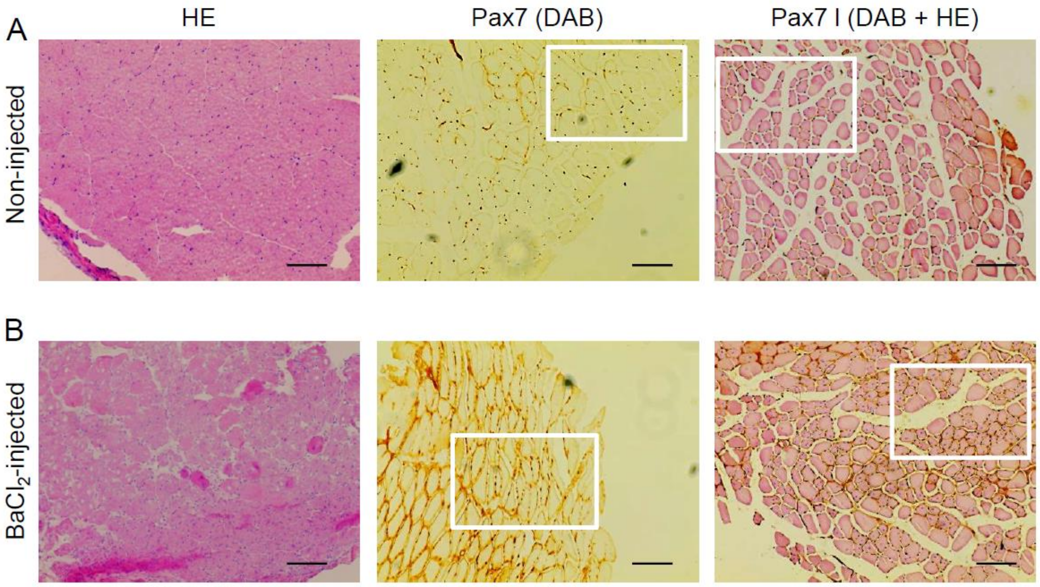

**Figure S8. Involvement of Pax7 in muscle regeneration. Related to Figure 6.** Cryosections from non-injected **(A)** and injected **(B)** TA muscle samples were subjected to Hematoxylin-Eosin (HE) and Pax7-specific immunostaining visualized with DAB reagent. Representative images from the aforementioned reactions were presented either alone (left and middle panels) or in combination (right panels). Scale bar is 100 µm. Specific areas from the representative images (indicated by squares) were further magnified (shown in Figure 6A-B) to represent different signal intensity for Pax7 staining and central nuclei in the injected samples.
